# Retest Reliability of Task-related fMRI BOLD Signals during Sequential Decision Making

**DOI:** 10.64898/2026.05.11.724283

**Authors:** Nicolas Leo Stege, Judit Pekar, Meyra Sophie Jackson, Filip Niemann, Miro Grundei, Ilinca-Maria Graur, Yiquan Shi, Shu-Chen Li

## Abstract

**Introduction:** Functional magnetic resonance imaging (fMRI) is widely used to study neural processes of behavior, but evaluations of test-retest reliability (TRR) of task-related blood-oxygen-level-dependent (BOLD) responses are scarce for many cognitive tasks. Such information is particularly important for longitudinal and intervention research. The ability to learn associations between choices and outcomes across decision stages is crucial for daily behavior. We assessed the measurement reliability of behavioral performance and fMRI BOLD signals during value-based sequential decision making to evaluate the TRR of task-relevant regions for future research on non-invasive brain stimulations.

**Methods:** Twenty healthy adults (22 to 40 years) completed two task-fMRI sessions, at least 2 weeks apart. During scanning, participants performed two variants of a three-stage Markov decision task with conditions varied in temporal contingency (immediate vs. delayed) and magnitude of choice outcomes (high vs. low). Both sessions were conducted under sham tDCS via a focal 3 × 1 montage targeting left dorsolateral prefrontal cortex (DLPFC). TRR was assessed using intraclass correlation coefficients (ICC) with a two-way mixed-effects consistency model for decision performance and task-related fMRI signals at voxel level and summarized in key regions defined by the extended Human Connectome Project atlas (HCPex).

**Results:** Decision performance was lower with delayed than immediate outcomes (*p* < 0.001). Higher outcome magnitude improved performance (*p* < 0.001). Decision performance increased across learning bins (*p* < 0.001). The behavioral TRR was in the moderate to good level (ICC(3,1) = 0.744 for accuracy; ICC(3,1) = 0.788 for reaction time). At the whole-brain level, contrasting brain activities in delayed with immediate condition revealed suprathreshold cluster peaks in several frontal-parietal (e.g., orbitofrontal, bilateral dorsolateral prefrontal, and medial parietal cortices) and striatal regions (e.g., bilateral putamen). Voxel-wise ICCs revealed variable but often moderate-to-good TRR across task-relevant regions, with stronger reliability in several striatal, orbitofrontal, and left dorsolateral prefrontal parcels, and more variable reliability across anterior cingulate and medial prefrontal parcels.

**Conclusion:** These results from a 2-session tDCS sham-sham stimulation study establish the validity of using the three-stage Markov decision task in future studies about intervention effects on the frontal-parietal-striatal network.

## 1 Introduction

Since the early 1990s, cognitive neuroscientists have used an array of measurement modalities to assess time-varying changes in blood-oxygen-level-dependent (BOLD) signals, which may reflect both spontaneous and task-induced changes in brain activities (Glover, 2011). Over the past three decades, functional Magnetic Resonance Imaging (fMRI) and associated methods have uncovered brain mechanisms underlying human behavior, ranging from perceptual, cognitive, emotional, and motivational processes. However, psychology and cognitive neuroscience have also been faced with a replication crisis (Huber et al., 2018; Lilienfeld & Strother, 2020) that is, in part, related to the low reliability of measured brain activities. Besides the issue of replicability in basic research, being able to reliably assess the effects of interventions is crucial for clinical research where therapeutic effects affect the health of individual patients (Elliott et al., 2020; Lachin, 2004).

To address these issues, measurement principles of assessing and achieving reliable measures need to be practiced. On the one hand, temporal variability in brain activities assessed either with fMRI or electroencephalogram (EEG) may not just reflect spontaneous physiological noise but could be systematically related to individual differences in cognitive performance and age (Garrett et al., 2013; Papenberg et al., 2013; Samanez-Larkin et al., 2010). On the other hand, such temporal variability and the individual difference therein make designing experimental tasks that yield sufficient measurement reliability quite challenging. Although the test-retest reliability (TRR) of fMRI task-related BOLD responses had been investigated in several earlier studies (e.g., Fliessbach et al., 2010; Plichta et al., 2012), it has not been a standard practice.

Meta-analytic evidence (Elliott et al., 2020) based on data from test-retest studies indicated that the TRR of several frequently used task-fMRI measures of BOLD responses assessed during clinically relevant psychological processes (e.g., cognitive control) is relatively poor, which compromises their clinical utility. More recently, utilizing datasets from the Human Connectome Project (https://www.humanconnectome.org), Wehrheim et al. (2024) analyzed the reliability in BOLD responses of seven tasks, covering important functional domains, including motor, emotional, reward, working memory, language, and social cognition. Their results showed moderate levels of TRR of task-related BOLD signals for most of these tasks. Whereas split-half reliability methods may yield higher reliability than TRR methods (Wehrheim et al., 2024), the latter is particularly important for intervention research which relies on repeated measurements in crossover designs.

This study investigated the TRR of BOLD responses during value-based sequential decisions, a cognitive ability crucial for the daily functioning of people across different ages, using the ICC. Individuals adapt their actions and decisions according to changing situational demands and behavioral goals. In many everyday contexts, decisions need to be made at successive stages, even though the information yielding crucial consequences may only become available much later than at the current stage when an initial decision (or action) is taken. Some of such sequential decisions can be formalized as a Markov process with the property that the choice made in each state not only determines this state’s immediate outcome but also influences the transition into upcoming decision states and their associated later outcomes (Tanaka et al., 2004; Eppinger et al., 2015).

Past evidence has consistently shown that brain regions such as the ventromedial PFC (vmPFC) and the striatum are active during valuation and reward processing (Rangel et al., 2008). Evidence from lesion studies, EEG recordings, and fMRI converges on activities in regions in the mid DLPFC (Brodmann areas 10, 45, and 9/46) as supporting the acquisition of abstract task rules, the representation of sequential relationships, and the integration of action–outcome contingencies across time for action planning and decision making (Badre & D’Esposito, 2007; Domenech & Koechlin, 2015; Badre & D’Esposito, 2009; Mansouri et al., 2020). Specifically, studies using sequential decision tasks with the Markov property (henceforth Markov task) showed that activities in several regions in the PFC and striatum implicate value-based learning across decision states to associate current choices with crucial outcomes that only occur at later states. In younger adults, BOLD responses in the left DLPFC tracked decision performance when value-based learning was prominent (Eppinger et al., 2015). Inhibiting this brain region using transcranial magnetic stimulation impaired younger adults’ decision performance (Wittkuhn et al., 2018). Moreover, adult age comparative (Eppinger et al., 2015) and cognitive intervention (Kang et al., 2024) studies revealed a lower recruitment of brain activities in several PFC regions in older adults or older adults who benefitted less from cognitive intervention. Of note, in comparison to younger adults, the performance of older adults was instead related to activities in the striatal regions, e.g. pallidum (Eppinger et al., 2015).

Given that the ability of sequential decision making is important for everyday behavior and it declines considerably during aging, non-invasive brain stimulation research has started to investigate stimulation protocols, such as transcranial direct current stimulation (tDCS), to enhance the efficiency of such decisions (e.g., Schommartz et al., 2021). Being part of a research consortium that systematically investigates effects of focalized tDCS on a broad range of cognitive and motor function using a double-blinded crossover design (https://www.memoslap.de/en/home/), this study assessed the TRR of a well-established Markov task with three decision stages (Eppinger et al., 2015; Kang et al., 2024; Wittkuhn et al., 2018) in several task-relevant regions in an initial sham-controlled intra-scanner crossover experiment with two task sessions. The main aim was to evaluate the measurement reliability of behavioral performance and task-fMRI activity during the 3-stage Markov task, which has not been examined before. Such information is important for establishing a basis for further investigations comparing intervention effects on sequential decision making over target and control stimulation sites.

To this end, we assessed intraclass correlation coefficient (ICC) of task-related measures at the behavioral and brain levels. The ICC is one of the most common methods for assessing TRR (Noble et al., 2019), which computes the degree of consistency between two measurement timepoints (Shrout & Fleiss, 1979). This concept is commonly denoted as the ICC(n,k), where “n” denotes the model used and “k” indicates the type of data averaging applied. There are multiple types of ICC models which can be employed. Here we used the ICC(3,1) that denotes a two-way mixed effects consistency model for examining whether the pattern or trend would be consistent. This model is more suitable for cases in which learning may affect the mean of measurement values in general, but the primary concern lies with the rank order of values across participants.

## 2 Materials and Methods

### 2.1 Study Overview and Experimental Design

The intra-scanner sham-controlled crossover study design and experimental procedure relevant for the current study are shown in Figure 1. After initial telephone screening during recruitment, as part of a larger study, each participant completed a baseline session involving an fMRI task-training session and a neuropsychological assessment. Participants also completed two fMRI task sessions (Task1, Task2), BOLD signals from the two sessions were used to evaluate the TRR of BOLD responses in task-relevant regions. The interval between the two task sessions was at least two weeks (M=20.45, SD=14.13 days).

**Figure 1.**
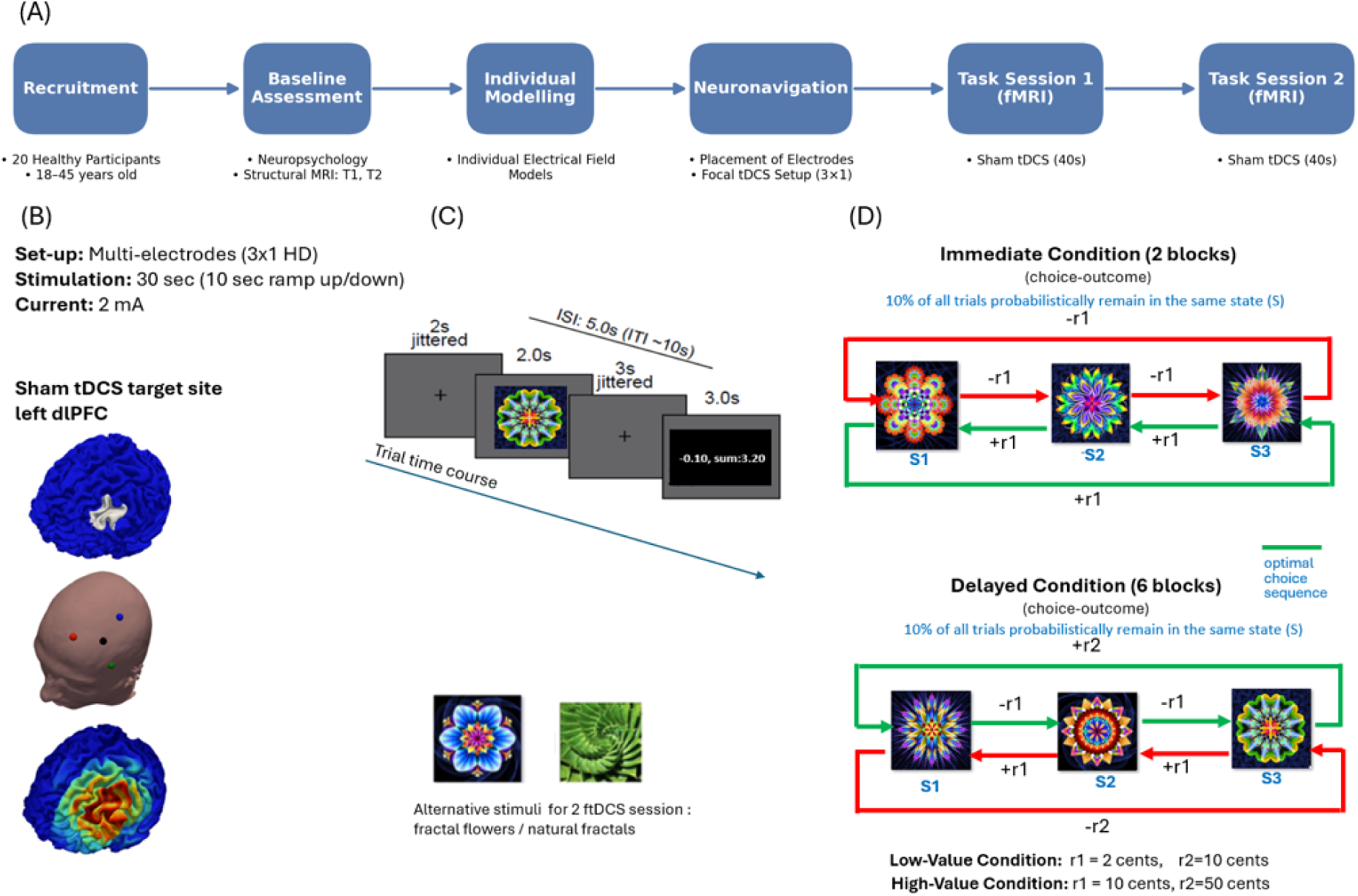
Study design, sham tDCS setup, and the 3-stage Markov task structure. Figure 1A: The different phases of the study are shown in upper panel horizontally as blue boxes progressing from left to right. Figure 1B: From top to bottom: target stimulation site over left dlPFC, Sham tDCS montage showing electrode placements, and current-flow simulation. Figure 1C: The trial structure of the 3-state Markov task, illustrating stimulus presentation, timing (in seconds), jitter intervals, and example stimuli (fractal flowers and natural fractals). Figure 1D: Structure of the three-state Markov decision task, illustrating sequential choice states under different outcome temporal contingencies (immediate vs. delayed) and outcome magnitudes.

#### 2.1.1 Participants

Twenty healthy adults (11 females) completed the experiment (aged 22–40 years, M=30.3, SD= 6.16). All participants had normal or corrected to normal vision and were right-handed assessed by the Edinburgh Handedness Inventory (Oldfield, 1971). All participants were fluent in German (C1 or first language) and did not report any history of neurological or psychological illness, alcoholism, drug use, or contraindications for MRI or tDCS (Antal et al., 2017). See Table 1 for details of demographics and baseline neuropsychological measures.

**Table 1.** Participant characteristics and baseline neuropsychological measures.

| Measure |  | M (SD) |
| --- | --- | --- |
| Age [years] |  | 30.3 (6.16) |
| Education [years] |  | 17.9 (2.39) |
| Beck Depression Inventory (BDI-II) |  | 3.93 (2.55) |
| Verbal Learning Memory (VLMT) [%] |  | 72.33 (12.14) |
| Spatial memory (RCFT) [%] |  | 74.19 (13.47) |
| Digit Span Test [n] | Digit forward | 7.45 (1.69) |
|  | Digit Backward | 6.7 (1.35) |
| Trail Making Test (TMT) [s] | A | 23.8 (6.51) |
|  | B | 61.38 (32.12) |
| Stroop Test [s] | Words | 29.01 (3.76) |
|  | Color | 47.26 (7.81) |
|  | Interference | 69.78 (12.72) |
| Regensburg Word Fluency Test (RWT) [n] | Phonemic | 16 (4.69) |
|  | Semantic | 25.8 (6.87) |
|  | Switch | 16.65 (3.18) |
| Multiple-Choice Vocabulary Test (MWT) [n] |  | 25.6 (5.04) |

*Descriptive statistics of participant demographics and the assessed neuropsychological measures.*

Before the MRI scans, participants underwent a neuropsychological assessment covering multiple cognitive domains. The battery included measures of verbal learning and memory assessed by the Verbal Learning and Memory Test (VLMT) (Müller et al., 1997), visuospatial memory measured with the Rey-Osterrieth Complex Figure Test (Zhang et al., 2021), short-term memory assessed by the digit span and reverse digit span (Woods et al., 2011), executive function measured with the Trail Making Test (Tombaugh, 2004) and the Stroop Color-Word Test (van der Elst et al., 2006), verbal fluency assessed by the Regensburg Verbal Fluency Test (Aschenbrenner et al., 2000), verbal intelligence and vocabulary knowledge (MWT; Lehrl, 1995), and fluid reasoning measured with a shortened version of the Ravens Test (Raven et al., 1998). Participants were also screened for depression using the Beck Depression Inventory (Kühner et al., 2007). Descriptive statistics for these measures are shown in Table 1. Participants also completed a baseline fMRI scan with a shortened training version of the Markov task. Based on their proportion of optimal choices in the last two delayed blocks during this baseline session, participants were stratified into higher performers (≥ 0.55; n = 7) and lower performers (< 0.55; n = 13).

#### 2.1.2 The 3-stage Markov Task

During fMRI acquisition, participants completed a first-order quasi-deterministic 3-stage Markov task (see Figure 1, lower middle and right panels). The task followed a first-order structure, meaning that state transitions were governed solely by the current state and the participant’s choice, and was quasi-deterministic, with transitions following a fixed mapping on most trials but including a 10% probability of stay trials. Within each trial, one of three sequential decision states, represented by an abstract visual stimulus, was shown to the participants who chose between two response options (left vs. right) at each state. At each state, a choice produces a positive or negative outcome with one of two magnitudes. The task was implemented with two conditions of temporal contingencies: an immediate condition in which the optimal option yielded a small gain at each of the three states (while the alternative yielded a small loss at each state); and a delayed condition in which the optimal decision policy required accepting small losses in earlier states to obtain a larger, later gain at the 3rd state (with the reversed pattern for the disadvantageous policy, i.e., accepting small gains that eventually lead to a large loss). Participants were instructed to maximize total gains by selecting actions that lead to overall gains and avoiding actions that lead to overall losses. They were informed that the gains and losses underlie different rules, which they could discover through the feedback provided after each response (Tanaka et al., 2004; Eppinger et al., 2015; Wittkuhn et al., 2018; Schommartz et al., 2021). In the current paradigm, two outcome magnitudes were also manipulated, with high-magnitude outcomes being 5-fold greater than their low-magnitude counterparts. The factor of temporal contingency (delayed vs. immediate) and outcome magnitude (high vs. low) were crossed, such that within each condition of temporal contingency (immediate vs. delayed), half the trials yielded low and the other half high outcome magnitudes. Two alternative versions of the tasks (with either images of fractal flowers or natural fractals as visual stimuli) were used in Task1 and Task2 sessions, so that carry-over effects can be minimized between task sessions. The assignment of the optimal response to left and right buttons was counterbalanced across participants. Each task session comprised 8 blocks of 30 trials each. The inter-trial interval (ITI) was approximately 10 s, ranging from 9 to 13 seconds. Jitter was applied to both the fixation cross duration (mean of 2 seconds) and the period after a response, before feedback was presented (mean of 3 seconds), with inter-stimulus intervals additionally jittered in 0.25-second steps (mean ISI of 5 seconds, see Figure 1C). If participants did not respond within a 2 s time window, a prompt appeared on the screen (“press 1 key!”), and the same stimulus was presented again. The same prompt was shown if participants pressed both buttons at the same time. The stimulus was repeated until a valid response was given. Trials with RT > 2000 ms, RT > 3 SDs above the mean reaction time, or more than one prompt appearance were excluded from subsequent analyses.

#### 2.1.3 MRI Acquisition

MRI data was acquired on a 3 Tesla Siemens Healthineers (MAGNETOM Prisma) scanner using a 64-channel head-neck coil in the Center of Cognitive Neuroscience Berlin at the Freie Universität Berlin. Task-based fMRI data were acquired using a 110 × 110 matrix with an in-plane spatial resolution of 2 × 2 mm² and 2-mm slice thickness with no interslice gap. Each participant underwent 2 task sessions that comprised 4 experimental runs that lasted about 45 min, each run averaging about 11 minutes in duration. The key acquisition parameters were: whole-brain; (T2-weighted gradient-echo EPI= 72 slices); interleaved order; repetition time (TR) = 1000 ms; echo time (TE) = 30.80 ms, flip angle (FA) = 60°, field of view (FOV) = 220 x 220 mm; and multiband acceleration factor = 6 and a structural scan (T1-weighted MPRAGE: 224 slices, TR = 2700, TE = 3.68 ms; 0.9 x 0.9 x 0.9 mm voxel). Each fMRI session lasted approximately 60 min, with phase encoding in the anterior-to-posterior (AP) direction.

#### 2.1.4 Sham focalized tDCS Setup

A focal 3 × 1 tDCS setup was used (see Figure 1B). Sham tDCS was administered during the two fMRI sessions using MRI-compatible equipment, specifically a multi-channel DC stimulator (DC-STIMULATOR MC, NeuroConn GmbH, Germany), circular conductive rubber electrodes (diameter = 2 cm), and cables. The sham stimulation protocol comprised a 10-s ramp-up and a ramp-down period, with 30 s of DC application at 2 mA in between. This protocol is designed to mimic the initial physical sensation typically associated with active tDCS without inducing sustained alterations in cortical excitability. The focal sham tDCS was administered over the left DLPFC, given previous evidence for its role in value-based sequential decision making (Eppinger et al., 2015; Wittkuhn et al., 2018). The coordinates of the electrode placements were selected for each participant in accordance with current flow and strength simulation results based on the participant’s structural brain images and guided by neuronavigation to maximize precision of individually tailored focalized tDCS (for details of the setup and stimulation protocol see Niemann et al. 2024).

### 2.2 Statistical Analysis

#### 2.2.1 Behavioral Data Analysis

To characterize the main behavioral effects of the task, behavioral performance (trial-level accuracy of making the optimal choice) during the Markov decision making was analyzed using a generalized linear mixed-effects model (GLMM) with a binomial distribution using a logit link function. Fixed effects included session (task1 vs. task2), outcome temporal contingency (delayed vs. immediate), outcome magnitude (high vs. low), learning bin (five consecutive bins of six trials), and interactions among these factors. Subject-specific random intercepts were also included.

To examine whether the effects of outcome temporal contingency on learning differed by initial performance level, a mixed ANOVA was conducted on decision performance averaged across sessions, with outcome temporal contingency (delayed vs. immediate), outcome magnitude (high vs. low), and learning bin as within-participant factors and performer group (higher vs. lower) as a between-participant factor. Greenhouse–Geisser corrections were applied where sphericity was violated. Follow-up ANOVAs were conducted within the delayed condition to characterize group-specific learning trajectories across bins.

#### 2.2.2 fMRI Data Analysis

The fMRI data of each participant were analyzed using SPM12 (r7771) implemented in MATLAB (R2021b). For the fMRI analysis the high performance computer at the ZEDAT, Freie Universität Berlin, was utilized (Bennett et al., 2020). Functional images were realigned using a two-pass procedure (register to mean) with 2nd-degree B-spline interpolation, and a mean functional image was generated, for each task session. The structural T1-weighted image from each session was coregistered to the mean functional image using normalized mutual information. The coregistered T1 was then segmented using SPM12’s unified segmentation, producing a bias-corrected image and a forward deformation field. Functional images were normalized to MNI space using the deformation field, resampled at 2×2×2 mm with 4th-degree B-spline interpolation. Finally, spatial smoothing was applied with an 8 mm FWHM Gaussian kernel.

For the first level analysis, a general linear model (GLM) was used to capture subject-specific task-related brain activity. Specifically, separate regressors were incorporated for the onsets of stimulus and feedback presentations for each combination of outcome temporal contingency (delayed vs. immediate) and outcome magnitude (high vs. low) and task block. Motion correction parameters (six realignment parameters) were included as nuisance regressors for each run. All experimental regressors were convolved with the canonical hemodynamic response function (HRF) as implemented in SPM12. A 128 s high-pass filter was applied.

First-level contrasts were computed separately for stimulus and feedback presentations. For the BOLD response analyses, contrast images were generated for each combination of temporal contingency (delayed vs. immediate) and outcome magnitude (high vs. low). For the primary ICC analysis, condition-specific beta estimates for the delayed condition were extracted separately for stimulus- and feedback-related activity. For an additional ICC analysis, a delayed > immediate contrast was also created.

At the second-level, separate flexible factorial models were created for stimulus-related and feedback-related contrasts. The models included eight conditions specific contrast images per participant (high-delay, low-delay, high-immediate, and low immediate for Task1 and Task2 sessions) and subject was included as a factor to account for repeated-measures. Statistical maps for initial voxel inclusion were thresholded at p < 0.001 (uncorrected) at the peak voxel level. Anatomical labels were obtained using the extended version of the Human Connectome Project multimodal parcellation atlas, HCPex. (Glasser et al., 2016; Huang et al., 2022).

Furthermore, in a separate analysis, the effects of performance level and age on brain activity were assessed. The performance vector (Perf), defined as decision accuracy in the delayed condition during fMRI sessions, was mean-centered (with higher value indicating higher performance) and entered the design matrix as a covariate (Perf_c). In addition, mean-centered age was included as a second covariate (Age_c, with larger value indicating older ages). Finally, a performance level by age interaction term was constructed as the product of the centered regressors (Perf_c x Age_c) and entered as a third covariate to assess whether age modulates the behavioral performance–BOLD association. Statistical maps for initial voxel inclusion were thresholded at p < 0.001 (uncorrected) at the peak voxel level.

#### 2.2.3 Test-Retest Reliability

The ICC analyses were conducted both for behavioral and the fMRI data using two-way mixed-effects consistency model, ICC(3,1), focusing only on data from the neurocognitively more demanding condition with delayed outcome temporal contingency, as the immediate condition served only as a control in this task. For behavioral performance, the ICC was performed for mean accuracy (choosing optimal option) and the mean reaction time (RT). The behavioral ICC analysis was computed in Python by implementing the two-way mixed-effects, consistency ICC using the Pingouin package (Vallat, 2018). To assess the TRR of the fMRI data, the Python-based Pyrelimri (Demidenko et al., 2024) tool was utilized to calculate ICCs based on whole-brain voxel-wise activation patterns. Following Pyrelimrís default method of handling of missing data (i.e., NaN values), the missing observations were replaced by the corresponding column means prior to ICC estimation. However, voxels which did not at least have valid values in both sessions for 10 participants were excluded. For the primary fMRI TRR analysis, ICCs were computed from the condition-specific delayed beta estimates rather than from the delayed > immediate contrast images. This choice was made for two reasons. Firstly because the delayed condition is the main condition of interest and secondly, as the delayed beta directly indexes the condition of interest, whereas contrast images behave as difference-score measures and may yield lower reliability by combining variance from two estimated condition responses, consistent with prior task-fMRI reliability work showing higher ICCs for beta coefficients than for contrasts (Heilicher et al., 2022). For each a priori parcel of interest previously implicated in value-based learning, the peak ICC voxel and its MNI coordinates, the 95% confidence interval at that peak, and the parcel-level mean ICC are reported.

## 3 Results

### 3.1 Behavioral Data

#### 3.1.1 Optimal Choice

The GLMM revealed that performance was higher in the immediate compared to the delayed condition (β = 0.60, SE = 0.16, z = 3.75, p < .001). Performance also improved across learning bins (all bins vs. bin 1: all p < .001), indicating successful learning over the course of the task. Critically, the effect of outcome temporal contingency interacted with learning bin, such that performance increased more steeply in the immediate condition, whereas learning in the delayed condition progressed more gradually (all interaction terms p < .001). A main effect of outcome magnitude was observed (β = 0.36, SE = 0.11, z = 3.20, p = .001), with higher outcome magnitude associated with better performance. No consistent interactions between outcome magnitude and learning bin were observed (all p > .60), indicating similar learning trajectories across magnitude conditions. Performance was also higher in the second session compared to the first (β = 0.18, SE = 0.05, z = 3.76, p < .001).

**Figure 2.**
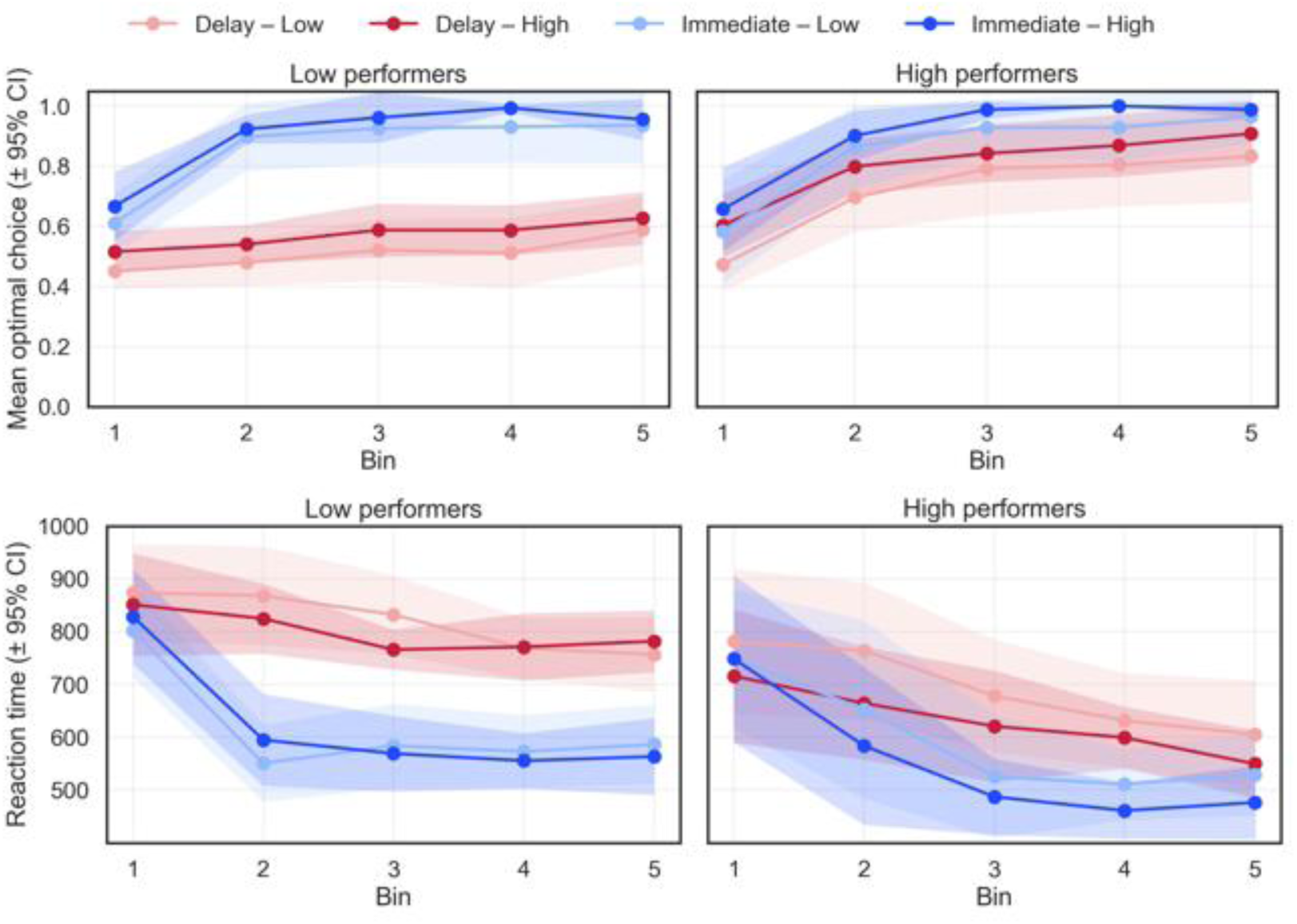
Behavioral results showing effects of outcome delay, outcome magnitude and individual difference in initial performance level. Optimal choice and RT across learning bins plotted separately for low and high outcome magnitude. Participants are stratified into high vs. low performers based on baseline task performance.

#### 3.1.2 Effects of Performance Group

To examine whether task performance differed as a function of initial performance level, we conducted a mixed analysis of variance (ANOVA) on optimal choices averaged across the two task sessions. Outcome temporal contingency (delayed vs. immediate), outcome magnitude (high vs. low), and learning bin (bins 1 to 5) were included as within-participant factors, and performer group (lower vs. higher performers) was included as a between-participant factor. Based on the initial performance assessed with a shortened version of the 3-stage Markov task at baseline, 13 participants were in the lower-performer group (proportion of choosing optimal options in the last two delayed blocks < 0.55) and 7 participants in the higher-performer group. Greenhouse-Geisser corrected p-values are reported for effects involving learning bin. To further characterize group-specific learning trajectories in the neurocognitively more demanding condition with delayed choice-outcome contingency, follow-up ANOVAs were conducted within the delayed condition only. These analyses tested whether lower and higher performers differed in delayed-condition learning across bins, and whether each group showed reliable changes in delayed-condition accuracy over time.

Regarding data on optimal choices, results of the ANOVA analysis showed that performer group interacted with outcome temporal contingency, F(1,18) = 14.59, p = .001, partial η² = .448, and with learning bin, F(4,72) = 5.37, p = .015, partial η² = .229. Importantly, the performer group × outcome temporal contingency × learning bin interaction was significant, F(4,72) = 5.96, p = .001, partial η² = .249, indicating that group differences in learning depended on outcome temporal contingency. This pattern reflected stronger group differences in the delayed condition than in the immediate condition.

Follow-up analyses within the delayed condition showed a significant performer group × learning bin interaction, F(4,72) = 8.08, p = .002, partial η² = .309. Higher performers showed a pronounced increase in delayed-condition accuracy across bins, F(4,24) = 49.67, p < .001, partial η² = .892. Lower performers also showed a significant bin effect, although this effect was smaller, F(4,48) = 4.61, p = .027, partial η² = .278. Outcome magnitude did not reliably interact with learning bin or performer group in the delayed-condition follow-up analysis.

The same mixed ANOVA structure was applied to reaction times (RTs). Responses became faster across learning bins, F(4,72) = 32.76, p < .001, partial η² = .645. Responses were also significantly faster in the immediate than delayed condition, F(1,18) = 39.32, p < .001, partial η² = .686. This difference varied by performer group, F(1,18) = 5.11, p = .036, partial η² = .221, reflecting a larger RT cost in the delayed condition among lower than higher performers. In addition, the change in RTs across learning bins differed by outcome temporal contingency, F(4,72) = 19.97, p < .001, partial η² = .526. RTs were also significantly faster for high- than low-magnitude outcomes, F(1,18) = 7.24, p = .015, partial η² = .287. However, RTs did not show a reliable performer group × outcome temporal contingency × learning bin interaction after Greenhouse-Geisser correction, F(4,72) = 2.78, p = .061, partial η² = .134.

### 3.2 Task fMRI Data

Considering previous evidence (Eppinger et al., 2015), we focused our analyses on brain activity patterns for the main effect of temporal contingency (contrasting delayed vs. immediate). An exploratory contrast comparing high outcome magnitude in the delayed condition vs. low outcome magnitude in the immediate condition was also examined. The statistical parametric maps were examined using no explicit masking and a voxel-wise peak threshold of p < 0.001 (uncorrected). Selected peaks are reported in the texts below and summarized in Figure 3, while all other activations that reached the statistical threshold are reported in Tables 2-4. Regions were defined and labeled using the HCPex Atlas (Glasser et al., 2016; Huang et al., 2022).

**Figure 3.**
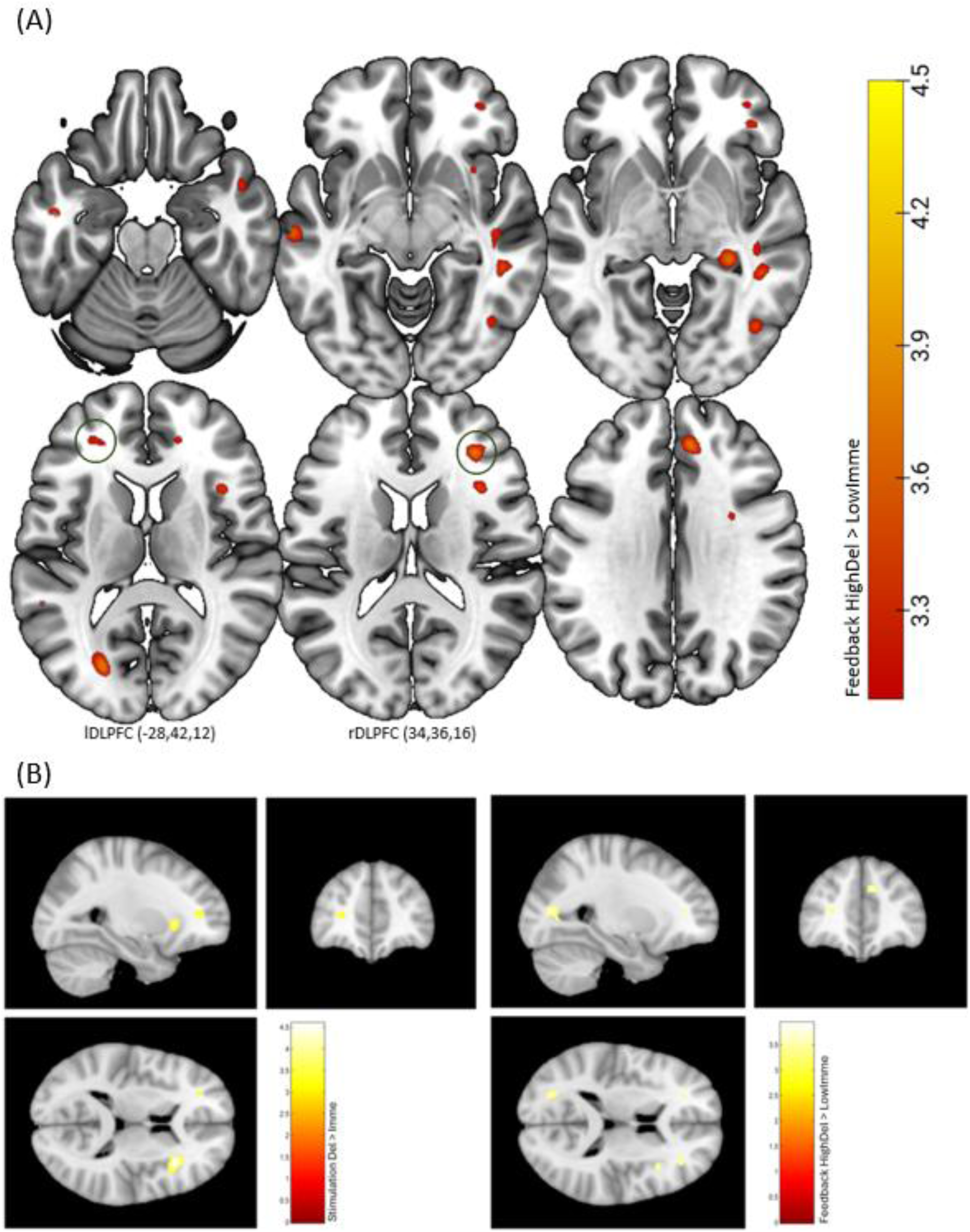
Whole-brain activation maps for the Delayed > Immediate and High_delayed > Low_immediate contrast at stimulus and feedback presentations. Figure 3a: Axial view slice montage of the High_Delayed > Low_Immediate contrast. Figure 3b: Left panel: SPM axial slice montages for the Delayed > Immediate contrast during stimulus presentation. Right panel: SPM axial slice montages for the High_Delayed > Low_Immediate contrast during feedback presentation.

**Table 2.**
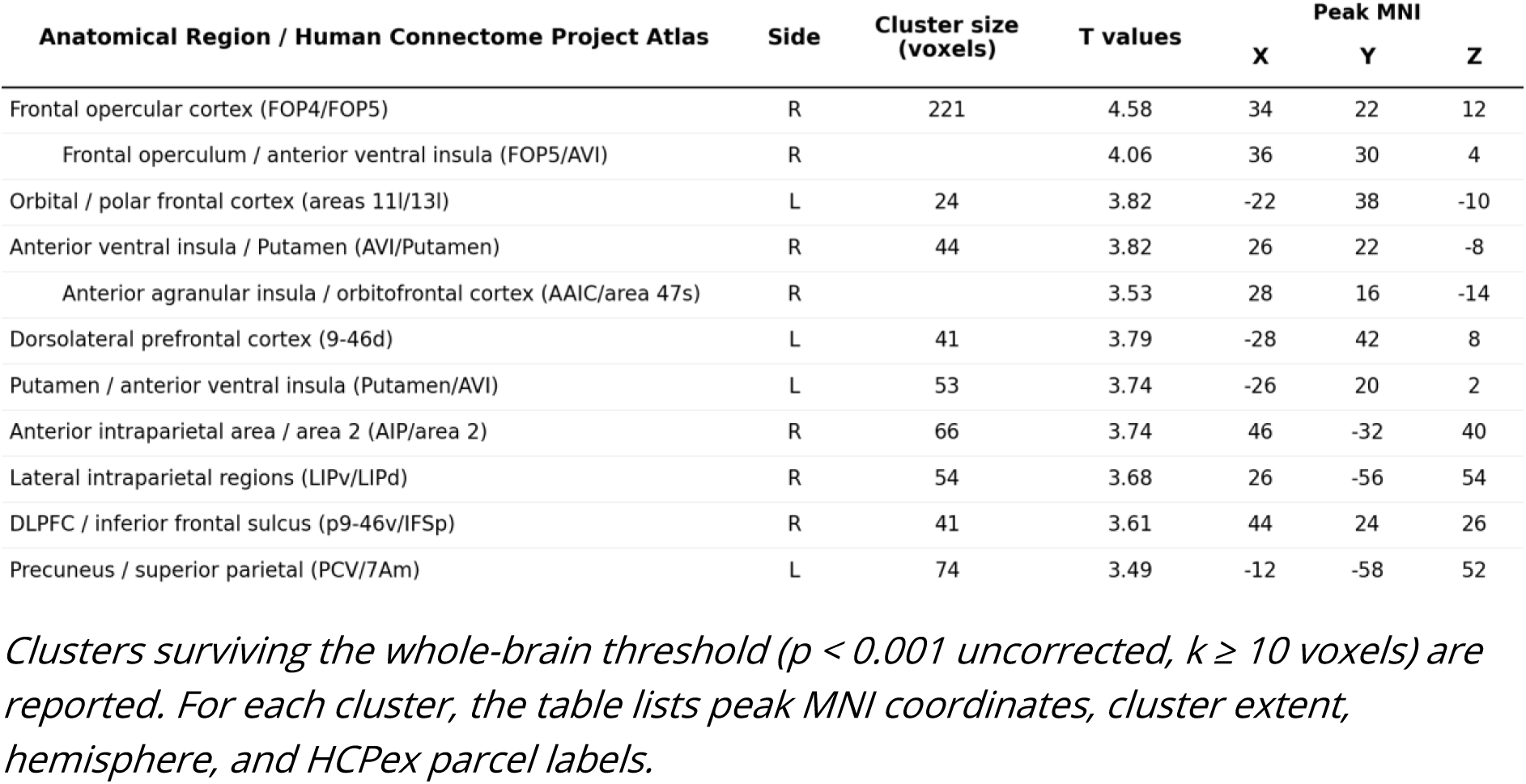
Whole-brain peaks showing greater activation in delayed compared to immediate condition during stimulus presentation.

#### 3.2.1 Effects at Stimulus Presentation

During stimulus presentation, the delayed vs. immediate contrast yielded higher activations in several regions in the frontal-striatal network (see results summarized in Table 2 and Figure 3 left panel). For instance, in voxels centered around the right insular-opercular cortex, with a peak in the right frontal operculum (MNI: 34, 22, 12; FOP4/FOP5) extending into right anterior ventral insular regions (MNI: 36, 30, 4; FOP5/AVI). Striatal involvement was also observed, primarily in the putamen (right: MNI 26, 22, −8; AVI/Putamen; left: MNI −26, 20, 2; Putamen/AVI), with putamen activity reaching threshold. In addition, significant clusters were present in bilateral DLPFC, with peaks in left DLPFC (MNI: −28, 42, 8; 9-46d/p47r) and right DLPFC (MNI: 44, 24, 26; p9-46v/IFSp).

#### 3.2.2 Effects at Feedback Presentation

During feedback presentation, the delayed vs. immediate contrast yielded higher activations in several frontal regions. For instance, in the right inferior frontal region, peaking at MNI coordinates 56, 32, 0 and encompassing Area 45 with extension into IFSa, indicating selective recruitment of right inferior frontal activation during delayed relative to immediate outcome (see results summarized in Table 3 for effects observed in other regions).

**Table 3.** Whole-brain peaks showing greater activation in delayed compared to immediate condition during feedback presentation.

| Anatomical Region / Human Connectome Project Atlas | Side | Cluster size (voxels) | T values | Peak MNI |  |  |
| --- | --- | --- | --- | --- | --- | --- |
|  |  |  |  | X | Y | Z |
| Inferior frontal cortex / inferior frontal sulcus (area 45/IFSa) | R | 51 | 3.61 | 56 | 32 | 0 |
| Area 47 lateral / area 45 (47l/45) | R |  | 3.35 | 50 | 30 | -8 |
| Para-insular area / superior temporal sulcus ventral anterior (PI/STSva) | L | 11 | 3.61 | -44 | -10 | -16 |
| Area TG dorsal / area TE1 anterior (TGd/TE1a) | L | 34 | 3.56 | -52 | 2 | -34 |
| Area STGa / area TG dorsal (STGa/TGd) | L |  | 3.46 | -52 | 10 | -26 |
| Auditory 5 complex / area TE1 middle (A5/TE1m) | L | 13 | 3.35 | -62 | -20 | -6 |

Clusters surviving the whole-brain threshold ( $p < 0.001$ uncorrected, $k \geq 10$ voxels) are reported. The table reports peak MNI coordinates, cluster extent, hemisphere, and HCPex parcel labels.

#### 3.2.3 Contrast Exploring Outcome Delay and Magnitude during Feedback

As shown in Figure 3 (right panel), greater activation associated with delayed outcomes of high magnitude relative to immediate outcomes of low magnitude during feedback presentation were observed in bilateral DLPFC, with a right peak in p9-46v/46 (MNI: 34, 36, 16) and a left peak in 9-46d extending into IFSa (MNI: −28, 42, 12). Inferior frontal peaks were also observed (MNI: 38, 48, -6) (see results summarized in Table 4 for effects of this contrast observed in other regions).

**Table 4.**
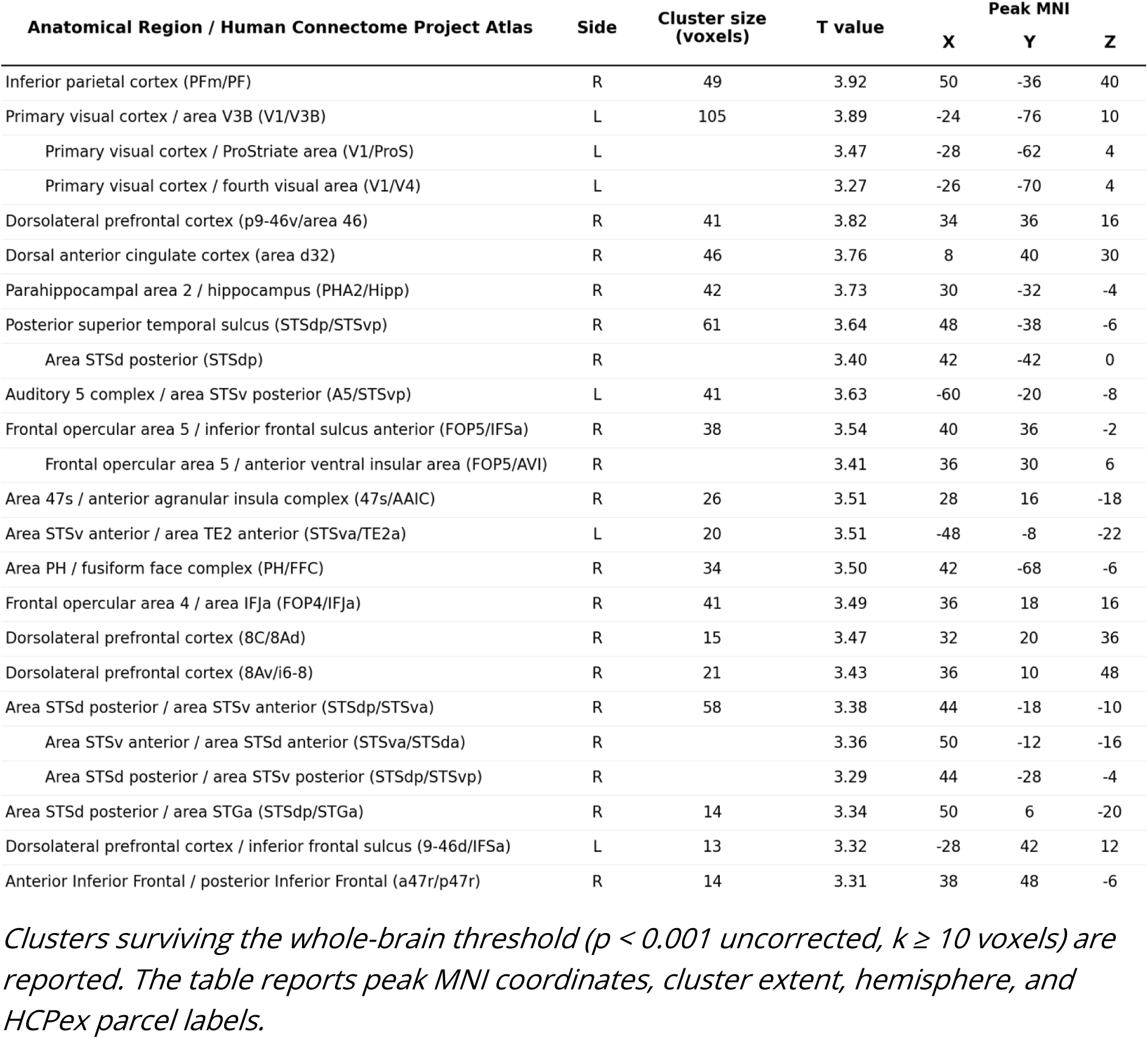
Whole-brain peaks for High_Delayed > Low_Immediate during Feedback presentation.

| Anatomical Region / Human Connectome Project Atlas | Side | Cluster size (voxels) | T value | Peak MNI |  |  |
| --- | --- | --- | --- | --- | --- | --- |
|  |  |  |  | X | Y | Z |
| Inferior parietal cortex (PFm/PF) | R | 49 | 3.92 | 50 | -36 | 40 |
| Primary visual cortex / area V3B (V1/V3B) | L | 105 | 3.89 | -24 | -76 | 10 |
| Primary visual cortex / ProStriate area (V1/ProS) | L |  | 3.47 | -28 | -62 | 4 |
| Primary visual cortex / fourth visual area (V1/V4) | L |  | 3.27 | -26 | -70 | 4 |
| Dorsolateral prefrontal cortex (p9-46v/area 46) | R | 41 | 3.82 | 34 | 36 | 16 |
| Dorsal anterior cingulate cortex (area d32) | R | 46 | 3.76 | 8 | 40 | 30 |
| Parahippocampal area 2 / hippocampus (PHA2/Hipp) | R | 42 | 3.73 | 30 | -32 | -4 |
| Posterior superior temporal sulcus (STSdp/STSvp) | R | 61 | 3.64 | 48 | -38 | -6 |
| Area STSd posterior (STSdp) | R |  | 3.40 | 42 | -42 | 0 |
| Auditory 5 complex / area STSv posterior (A5/STSvp) | L | 41 | 3.63 | -60 | -20 | -8 |
| Frontal opercular area 5 / inferior frontal sulcus anterior (FOP5/IFSa) | R | 38 | 3.54 | 40 | 36 | -2 |
| Frontal opercular area 5 / anterior ventral insular area (FOP5/AVI) | R |  | 3.41 | 36 | 30 | 6 |
| Area 47s / anterior agranular insula complex (47s/AAIC) | R | 26 | 3.51 | 28 | 16 | -18 |
| Area STSv anterior / area TE2 anterior (STSva/TE2a) | L | 20 | 3.51 | -48 | -8 | -22 |
| Area PH / fusiform face complex (PH/FFC) | R | 34 | 3.50 | 42 | -68 | -6 |
| Frontal opercular area 4 / area IFJa (FOP4/IFJa) | R | 41 | 3.49 | 36 | 18 | 16 |
| Dorsolateral prefrontal cortex (8C/8Ad) | R | 15 | 3.47 | 32 | 20 | 36 |
| Dorsolateral prefrontal cortex (8Av/i6-8) | R | 21 | 3.43 | 36 | 10 | 48 |
| Area STSd posterior / area STSv anterior (STSdp/STSva) | R | 58 | 3.38 | 44 | -18 | -10 |
| Area STSv anterior / area STSd anterior (STSva/STSda) | R |  | 3.36 | 50 | -12 | -16 |
| Area STSd posterior / area STSv posterior (STSdp/STSvp) | R |  | 3.29 | 44 | -28 | -4 |
| Area STSd posterior / area STGa (STSdp/STGa) | R | 14 | 3.34 | 50 | 6 | -20 |
| Dorsolateral prefrontal cortex / inferior frontal sulcus (9-46d/IFSa) | L | 13 | 3.32 | -28 | 42 | 12 |
| Anterior Inferior Frontal / posterior Inferior Frontal (a47r/p47r) | R | 14 | 3.31 | 38 | 48 | -6 |

*Clusters surviving the whole-brain threshold ( $p < 0.001$ uncorrected, $k \geq 10$ voxels) are reported. The table reports peak MNI coordinates, cluster extent, hemisphere, and HCPex parcel labels.*

#### 3.2.4 Effects of Performance x Age Interaction

The Performance × Age interaction was examined separately for stimulus and feedback presentation, and for the Task1 and Task2 sessions. A significant positive interaction was observed in bilateral DLPFC and frontoparietal regions, indicating that performance-related increases in BOLD responses were more pronounced in older participants in our sample.

During stimulus presentation, the most prominent peaks were observed in bilateral DLPFC in the first session (right 8Ad/46, MNI: 28, 36, 40; left 8Ad, MNI: -26, 34, 40), together with an OFC peak in the first session (11l/13l, MNI: 22, 42, - 10; see Figure 4, left panel). In the second session, stimulus-related effects were more circumscribed and centred on medial frontal/cingulate cortex (p32pr/24dv, MNI: -18, 6, 38), with an additional right medial temporal peak (TF/Hippocampus, MNI: 42, -22, -18).

**Figure 4.**
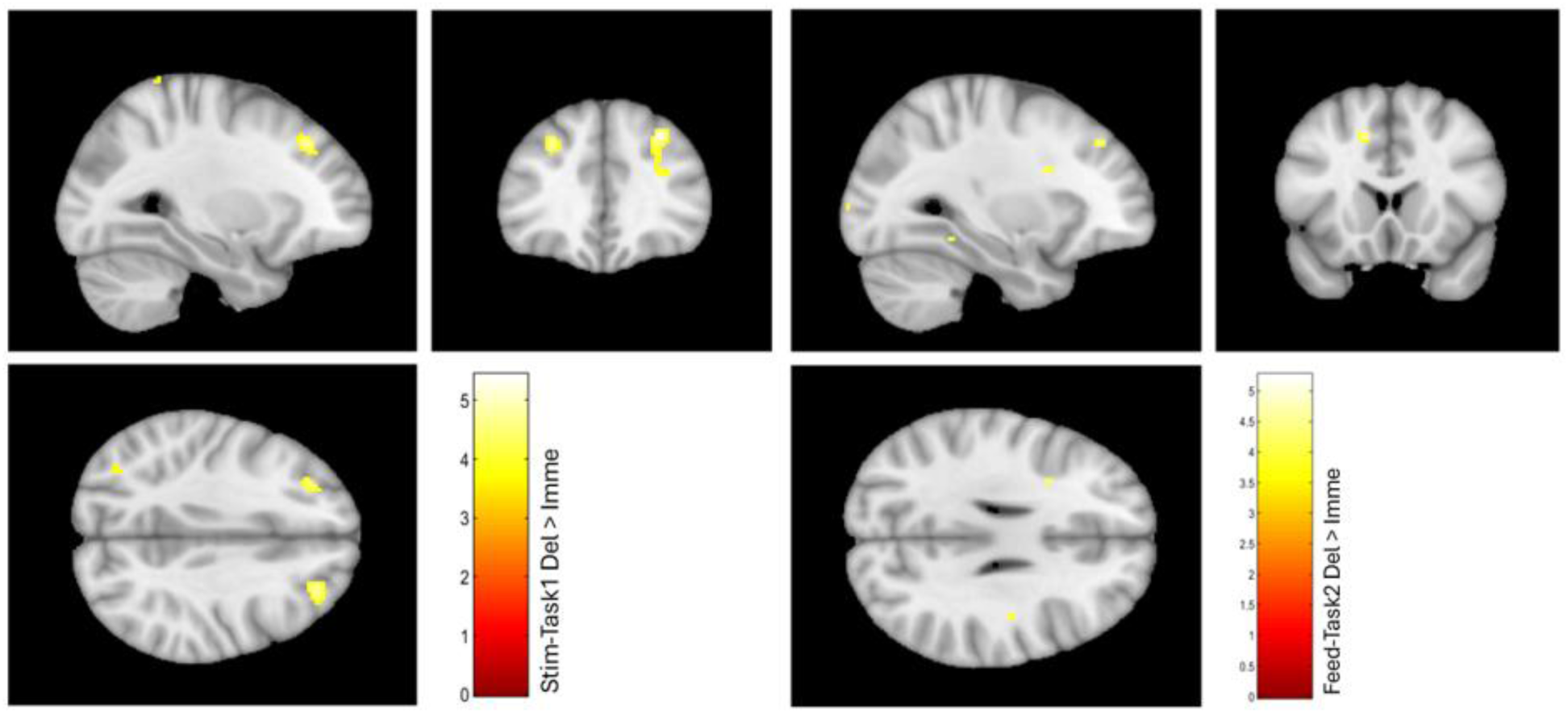
Whole-brain regression maps for the Performance × Age interaction. Group-level regression results showing selected task-related activities associated with the Performance × Age covariate, displayed separately for stimulus-related activity in Task1 (left) and feedback-related activity in Task2.

During feedback presentation, the interaction again involved positive associations with the BOLD response of the DLPFC, with bilateral DLPFC peaks in the first session (right 46/9-46d, MNI: 30, 32, 22; left 9-46d/46, MNI: -26, 38, 20) and left DLPFC/medial frontal peaks in the second session (8Ad/p32pr, MNI: -20, 26, 34; p32pr/24dv, MNI: -18, 6, 38). Additional peaks, including parietal, temporal, premotor, cingulate, and striatal regions, are summarized in Figure 4 (right panel) and Table 5.

**Table 5.** Whole-brain peak voxels showing positive Performance × Age interaction effects on task-evoked BOLD activity.

| Event | Task | HCPex Parcel | Region | Side | Peak MNI |  |  |
| --- | --- | --- | --- | --- | --- | --- | --- |
|  |  |  |  |  | X | Y | Z |
| Stimulus | Task 1 | 8Ad/46 | DLPFC | R | 28 | 36 | 40 |
| Stimulus | Task 1 | 8Ad | DLPFC | L | -26 | 34 | 40 |
| Stimulus | Task 1 | 11l/13l | OFC | R | 22 | 42 | -10 |
| Stimulus | Task 1 | 6mp/4 | Motor cortex | L | -10 | -16 | 76 |
| Stimulus | Task 1 | 5L | Superior parietal | R | 12 | -42 | 72 |
| Stimulus | Task 1 | 4/3b | Motor/somatosensory | R | 10 | -24 | 76 |
| Stimulus | Task 1 | AIP | Parietal cortex | R | 36 | -42 | 50 |
| Stimulus | Task 1 | 7AL/5L | Posterior parietal cortex | L | -26 | -48 | 74 |
| Stimulus | Task 2 | p32pr/24dv | Medial frontal/cingulate | L | -18 | 6 | 38 |
| Stimulus | Task 2 | TF/Hippocampus | Medial temporal cortex | R | 42 | -22 | -18 |
| Stimulus | Task 2 | 8BL/8AD | DLPFC | R | 20 | 46 | 44 |
| Feedback | Task 1 | TE2a/TE2p | Lateral temporal cortex | R | 54 | -36 | -18 |
| Feedback | Task 1 | 46/9-46d | DLPFC | R | 30 | 32 | 22 |
| Feedback | Task 1 | 7AL/5L | Superior parietal cortex | L | -24 | -46 | 72 |
| Feedback | Task 1 | Pol2/TA2 | Temporal pole/auditory cortex | R | 44 | -2 | -6 |
| Feedback | Task 1 | 9-46d/46 | DLPFC | L | -26 | 38 | 20 |
| Feedback | Task 2 | 8Ad/p32pr | DLPFC/medial frontal | L | -20 | 26 | 34 |
| Feedback | Task 2 | p32pr/24dv | Medial frontal/cingulate | L | -18 | 6 | 38 |
| Feedback | Task 2 | 6a | Premotor cortex | L | -28 | 8 | 54 |
| Feedback | Task 2 | Caudate | Striatum | L | -22 | -28 | 26 |
| Feedback | Task 2 | VMV2/VMV1 | Ventral visual cortex | L | -30 | -46 | 4 |
| Feedback | Task 2 | 31pv/RSC | Posterior cingulate/retrosplenial | L | -16 | -44 | 24 |
| Feedback | Task 2 | a32pr | Medial frontal cortex | R | 18 | 24 | 22 |

*Whole-brain peak voxels showing positive Performance × Age interaction effects on task-evoked BOLD activity during stimulus- and feedback-related processing in two task sessions.*

### 3.3 Test-Retest Reliability

As an overview, ICC values range from 0 to 1, with interpretative guidelines suggesting that values below 0.50 are indicative of poor test-retest reliability, values between 0.50 and 0.75 are judged as moderate reliability, values between 0.75 and 0.90 are considered to reflect good reliability, and values above 0.90 are regarded as excellent reliability (Koo & Li, 2016).

#### 3.3.1 Behavioral TRR Results

To quantify the reliability across the two task versions, we computed a two-way mixed-effects, single-measure ICC (3,1) using each participant’s (n = 20) mean score per task (averaged across time bins). Accuracy (proportion of choosing optimal options) showed an ICC of 0.744 (95% CI [0.460, 0.890]), reflecting moderate point estimate consistency in performance between the two task sessions. The reaction times yielded an ICC of 0.788 (95% CI [0.540, 0.910]), indicating good point estimate reliability. Overall, participants’ performance was notably consistent across sessions, supporting the robustness of these behavioral measures across task variants and despite learning effects.

#### 3.3.2 fMRI Results – Stimulus Presentation

To assess TRR of BOLD responses during stimulus presentation, ICC maps were computed based on the condition-specific beta estimates, relative to baseline, for the delayed condition and overlaid with the HCPex atlas. For parcels in a priori defined regions of interest previously shown to be involved in value-based learning, the peak ICC voxel and its MNI coordinate were extracted. Only voxels with at least 10 participants contributing finite paired values were retained. ICC results for the main effect contrast, delayed vs. immediate, can be found in the supplementary material, along with results computed without the finite paired value voxel criterion.

Based on ICC point estimates, left DLPFC parcels demonstrated predominantly moderate peak ICCs, with the highest point estimate reliability found in 8BL, MNI: −8, 42, 54, ICC = 0.784, Area 46, MNI: −36, 36, 30, ICC = 0.740, and 8C, MNI: −40, 0, 38, ICC = 0.721. ICC values for all other key regions are presented in Table 6 and summarized as an overview in Figure 5, left panel.

**Table 6.** Peak ICC values of key regions defined by the HCPex atlas for task fMRI BOLD responses during Stimulus presentation.

| Parcel | Cortical Division | Side | Mean ICC | Peak ICC | 95% CI | Peak MNI (x, y, z) |
| --- | --- | --- | --- | --- | --- | --- |
| **DLPFC** |  |  |  |  |  |  |
| 46 | Dorsolateral_Prefrontal | L | 0.3713 | 0.7401 | [0.4521, 0.8884] | (-36.0, 36.0, 30.0) |
| 8Ad | Dorsolateral_Prefrontal | L | 0.0786 | 0.5323 | [0.1292, 0.7844] | (-30.0, 22.0, 46.0) |
| 8Av | Dorsolateral_Prefrontal | L | 0.2743 | 0.5682 | [0.1796, 0.8035] | (-44.0, 26.0, 42.0) |
| 8BL | Dorsolateral_Prefrontal | L | 0.3785 | 0.7841* | [0.5316, 0.9086] | (-8.0, 42.0, 54.0) |
| 8C | Dorsolateral_Prefrontal | L | 0.4968 | 0.7217 | [0.4200, 0.8797] | (-40.0, 0.0, 38.0) |
| 9-46d | Dorsolateral_Prefrontal | L | 0.1827 | 0.6508 | [0.3035, 0.8455] | (-26.0, 50.0, 6.0) |
| 9a | Dorsolateral_Prefrontal | L | 0.1189 | 0.6617 | [0.3207, 0.8509] | (-14.0, 54.0, 28.0) |
| 9p | Dorsolateral_Prefrontal | L | 0.3347 | 0.6640 | [0.3244, 0.8520] | (-12.0, 52.0, 30.0) |
| a9-46v | Dorsolateral_Prefrontal | L | 0.3027 | 0.6039 | [0.2316, 0.8219] | (-40.0, 60.0, 2.0) |
| i6-8 | Dorsolateral_Prefrontal | L | 0.3493 | 0.5822 | [0.1997, 0.8107] | (-24.0, 6.0, 60.0) |
| s6-8 | Dorsolateral_Prefrontal | L | 0.3321 | 0.6761 | [0.3439, 0.8579] | (-22.0, 22.0, 62.0) |
| SFL | Dorsolateral_Prefrontal | L | 0.2338 | 0.6280 | [0.2680, 0.8341] | (-8.0, 2.0, 70.0) |
| **vmPFC/OFC and ACC/mPFC** |  |  |  |  |  |  |
| 10r | AntCing_MedPFC | L | 0.3420 | 0.6775 | [0.3462, 0.8586] | (-6.0, 60.0, -10.0) |
| 10v | AntCing_MedPFC | L | 0.4687 | 0.8090* | [0.5789, 0.9198] | (-6.0, 54.0, -18.0) |
| 25 | AntCing_MedPFC | L | 0.2594 | 0.6472 | [0.2978, 0.8437] | (-2.0, 14.0, -14.0) |
| a24 | AntCing_MedPFC | L | 0.4341 | 0.6628 | [0.3225, 0.8514] | (0.0, 44.0, -6.0) |
| a32pr | AntCing_MedPFC | L | 0.1965 | 0.3415 | [-0.1072, 0.6746] | (-8.0, 24.0, 36.0) |
| d32 | AntCing_MedPFC | L | 0.2276 | 0.5152 | [0.1060, 0.7752] | (-6.0, 50.0, 32.0) |
| pOFC | AntCing_MedPFC | L | 0.2698 | 0.5809 | [0.1978, 0.8101] | (-12.0, 16.0, -14.0) |
| s32 | AntCing_MedPFC | L | 0.1395 | 0.4362 | [0.0041, 0.7310] | (-8.0, 40.0, -8.0) |
| 11l | OrbPolaFrontal | L | 0.4666 | 0.8825* | [0.7275, 0.9518] | (-30.0, 48.0, -14.0) |
| 13l | OrbPolaFrontal | L | 0.2981 | 0.7099 | [0.4000, 0.8742] | (-16.0, 42.0, -18.0) |
| 47m | OrbPolaFrontal | L | 0.2330 | 0.5134 | [0.1036, 0.7742] | (-28.0, 36.0, -14.0) |
| OFC | OrbPolaFrontal | L | 0.3547 | 0.8084* | [0.5777, 0.9195] | (-10.0, 50.0, -20.0) |
| a47r | Inferior_Frontal | L | 0.2456 | 0.8212* | [0.6026, 0.9252] | (-34.0, 48.0, -12.0) |
| p47r | Inferior_Frontal | L | 0.2801 | 0.5868 | [0.2063, 0.8131] | (-46.0, 40.0, 6.0) |
| 10r | AntCing_MedPFC | R | 0.4261 | 0.7172 | [0.4123, 0.8776] | (8.0, 60.0, -6.0) |
| 10v | AntCing_MedPFC | R | 0.4181 | 0.7482 | [0.4663, 0.8921] | (4.0, 52.0, -20.0) |
| 25 | AntCing_MedPFC | R | 0.1860 | 0.6741 | [0.3406, 0.8569] | (6.0, 18.0, -18.0) |
| a24 | AntCing_MedPFC | R | 0.5255 | 0.7694* | [0.5045, 0.9019] | (6.0, 38.0, -2.0) |
| a32pr | AntCing_MedPFC | R | 0.3193 | 0.5706 | [0.1829, 0.8047] | (10.0, 22.0, 40.0) |
| d32 | AntCing_MedPFC | R | 0.1848 | 0.4326 | [-0.0003, 0.7289] | (6.0, 40.0, 34.0) |
| pOFC | AntCing_MedPFC | R | 0.2109 | 0.7819* | [0.5276, 0.9076] | (22.0, 10.0, -22.0) |
| s32 | AntCing_MedPFC | R | 0.2936 | 0.5427 | [0.1436, 0.7900] | (2.0, 42.0, -10.0) |
| 11l | OrbPolaFrontal | R | 0.4250 | 0.8398* | [0.6393, 0.9333] | (18.0, 40.0, -20.0) |
| 13l | OrbPolaFrontal | R | 0.3651 | 0.7257 | [0.4270, 0.8816] | (20.0, 42.0, -18.0) |
| 47m | OrbPolaFrontal | R | 0.4250 | 0.7185 | [0.4146, 0.8782] | (30.0, 38.0, -6.0) |
| OFC | OrbPolaFrontal | R | 0.3182 | 0.7783* | [0.5209, 0.9059] | (14.0, 40.0, -20.0) |
| a47r | Inferior_Frontal | R | 0.2527 | 0.7729* | [0.5110, 0.9035] | (28.0, 52.0, -4.0) |
| p47r | Inferior_Frontal | R | 0.4474 | 0.6868 | [0.3614, 0.8631] | (46.0, 40.0, -6.0) |
| **Striatum** |  |  |  |  |  |  |
| Putam | Subcortical | L | 0.3016 | 0.7524 | [0.4739, 0.8941] | (-28.0, 8.0, 8.0) |
| Caud | Subcortical | L | 0.3396 | 0.7614 | [0.4900, 0.8982] | (-8.0, 10.0, 16.0) |
| NAc | Subcortical | L | -0.1580 | 0.2851 | [-0.1685, 0.6391] | (-12.0, 8.0, -8.0) |
| Putam | Subcortical | R | 0.3692 | 0.7795* | [0.5232, 0.9065] | (28.0, 16.0, -4.0) |
| Caud | Subcortical | R | 0.4296 | 0.8258* | [0.6116, 0.9272] | (14.0, 6.0, 22.0) |
| NAc | Subcortical | R | 0.0490 | 0.6019 | [0.2287, 0.8209] | (14.0, 8.0, -8.0) |

For parcels in each a priori ROI, the table summarizes the location of the most reliable voxel during stimulus processing, together with its peak intraclass correlation coefficient (ICC) and MNI coordinates for BOLD responses in the delayed condition. Asterisks indicate the reliability level of the peak ICC based on the lower bound of the 95% confidence interval: \* moderate reliability ( $> 0.50$ ), \*\* good reliability ( $> 0.75$ ), \*\*\* excellent reliability ( $> 0.90$ ).

**Figure 5.**
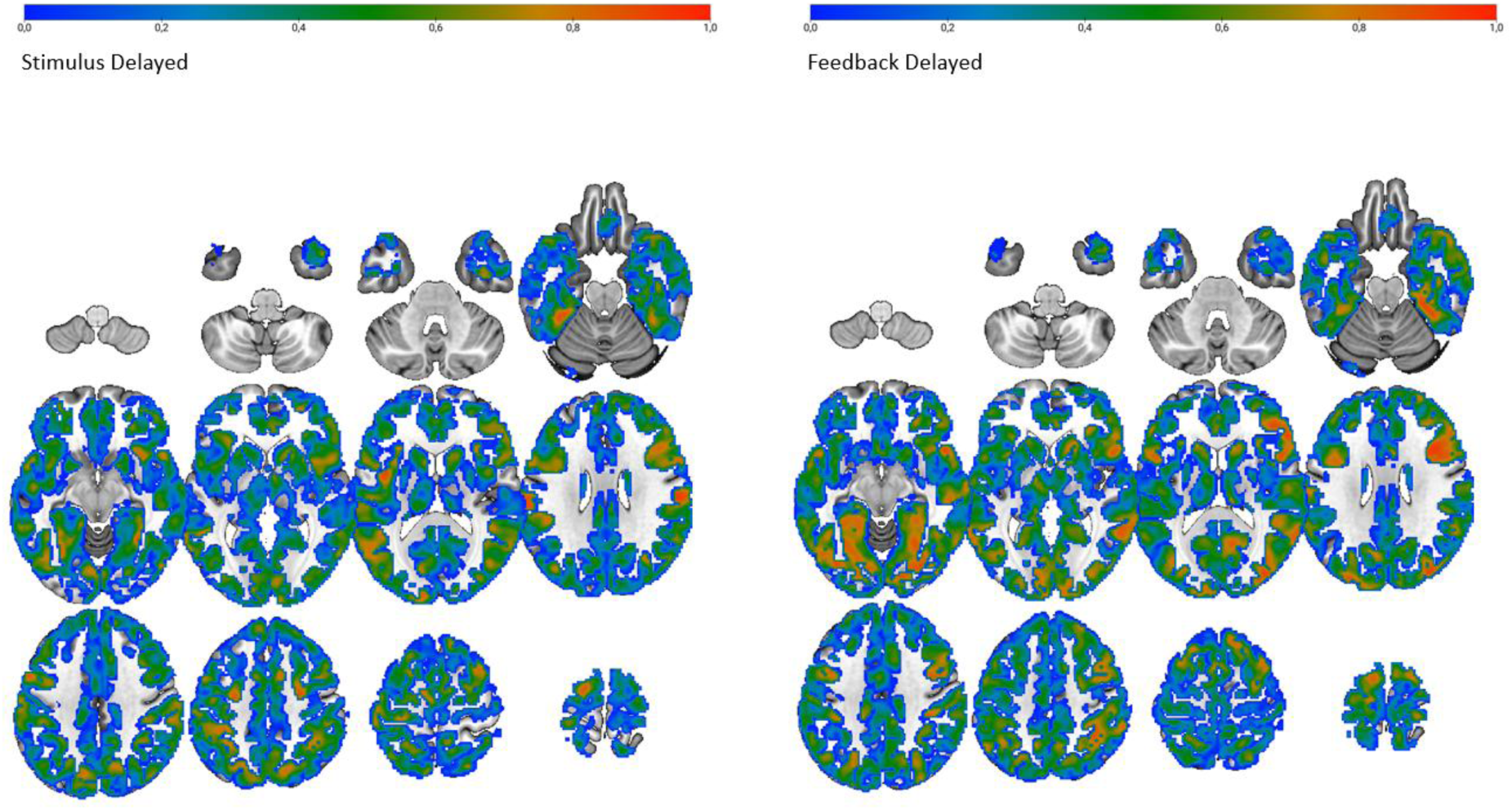
Whole-brain voxel-wise heat map for TRR of condition-specific activation for BOLD responses in the delayed condition. Whole-brain voxel-wise ICC map computed for the delayed condition specific regressor, for stimulus-related activity (left) and feedback-related activity (right). Only regions that fall under the HCPex atlas are shown.

Striatal parcels showed generally moderate-to-good reliability for caudate and putamen, with more variable reliability in nucleus accumbens. In the left hemisphere, putamen, MNI: −28, 8, 8, ICC = 0.752, and caudate, MNI: −8, 10, 16, ICC = 0.761, demonstrated good peak reliability, while nucleus accumbens showed poor reliability, MNI: −12, 8, −8, ICC = 0.285. In the right hemisphere, reliability was similarly high for caudate, MNI: 14, 6, 22, ICC = 0.825, and putamen, MNI: 28, 16, −4, ICC = 0.779, and moderate for nucleus accumbens, MNI: 14, 8, −8, ICC = 0.601.

Of the 12 left DLPFC parcels examined, one parcel, 8BL, had a 95% CI lower bound exceeding 0.50, indicating that at least moderate reliability can be statistically supported for stimulus-related activation in this region. Among the extended regions, confirmed reliability differed across cortical divisions during stimulus presentation. In ACC/mPFC parcels, 1 of 8 left-hemisphere parcels and 2 of 8 right-hemisphere parcels exceeded lower CI bound of 0.50. In OFC/vmPFC parcels, 3 of 6 parcels in the left hemisphere and 3 of 6 parcels in the right hemisphere exceeded this threshold. Striatal parcels showed limited confirmed reliability, with 2 of 6 parcels exceeding the 0.50 lower-bound threshold, specifically right putamen and right caudate.

#### 3.3.3 fMRI Results – Feedback Presentation

The same procedure as for stimulus presentation was used to compute the ICC of the BOLD activity during feedback presentation.

Based on ICC point estimates, left DLPFC parcels showed predominantly moderate-to-good voxel-wise ICC values at their peak locations. The highest reliability was observed in left Area 8C, MNI: −40, 4, 34, ICC = 0.899, followed by left SFL, MNI: −10, 16, 72, ICC = 0.817, left i6-8, MNI: −30, 12, 60, ICC = 0.800, left 8Av, MNI: −30, 14, 60, ICC = 0.791, left a9-46v, MNI: −48, 46, 14, ICC = 0.785, and left Area 46, MNI: −42, 46, 24, ICC = 0.779. The lowest peak reliability within the set was observed in left 8Ad, MNI: −24, 28, 40, ICC = 0.609, though it still indicated moderate reliability.

Striatal regions showed moderate-to-good reliability across hemispheres, with stronger reliability in caudate and putamen than in nucleus accumbens. In the left hemisphere, caudate, MNI: −16, 18, 16, ICC = 0.833, and putamen, MNI: −24, 18, −2, ICC = 0.805, achieved good peak ICCs. In the right hemisphere, comparable reliability was observed in caudate, MNI: 18, 16, 4, ICC = 0.833, and putamen, MNI: 28, −12, 10, ICC = 0.831, while nucleus accumbens showed moderate reliability, MNI: 14, 8, −8, ICC = 0.632. ICC values for all other key regions are presented in Table 7 and summarized as an overview in Figure 5, right panel.

**Table 7.** Peak ICC values of key regions defined by the HCPex atlas during Feedback presentation.

| Parcel | Cortical Division | Side | Mean ICC | Peak ICC | 95% CI | Peak MNI (x, y, z) |
| --- | --- | --- | --- | --- | --- | --- |
| **DLPFC** |  |  |  |  |  |  |
| 46 | Dorsolateral_Prefrontal | L | 0.3131 | 0.7798* | [0.5237, 0.9066] | (-42.0, 46.0, 24.0) |
| 8Ad | Dorsolateral_Prefrontal | L | 0.3311 | 0.6095 | [0.2400, 0.8248] | (-24.0, 28.0, 40.0) |
| 8Av | Dorsolateral_Prefrontal | L | 0.3353 | 0.7910* | [0.5446, 0.9117] | (-30.0, 14.0, 60.0) |
| 8BL | Dorsolateral_Prefrontal | L | 0.3880 | 0.7403 | [0.4526, 0.8885] | (-8.0, 38.0, 56.0) |
| 8C | Dorsolateral_Prefrontal | L | 0.3564 | 0.8990** | [0.7630, 0.9588] | (-40.0, 4.0, 34.0) |
| 9-46d | Dorsolateral_Prefrontal | L | 0.3749 | 0.7509 | [0.4712, 0.8934] | (-38.0, 50.0, 22.0) |
| 9a | Dorsolateral_Prefrontal | L | 0.2425 | 0.7397 | [0.4514, 0.8882] | (-26.0, 54.0, 32.0) |
| 9p | Dorsolateral_Prefrontal | L | 0.3633 | 0.7440 | [0.4589, 0.8902] | (-26.0, 50.0, 34.0) |
| a9-46v | Dorsolateral_Prefrontal | L | 0.2525 | 0.7856* | [0.5345, 0.9093] | (-48.0, 46.0, 14.0) |
| i6-8 | Dorsolateral_Prefrontal | L | 0.4638 | 0.8001* | [0.5618, 0.9158] | (-30.0, 12.0, 60.0) |
| s6-8 | Dorsolateral_Prefrontal | L | 0.4950 | 0.7759* | [0.5166, 0.9049] | (-20.0, 22.0, 62.0) |
| SFL | Dorsolateral_Prefrontal | L | 0.5271 | 0.8176* | [0.5955, 0.9236] | (-10.0, 16.0, 72.0) |
| **vmPFC/OFC and ACC/mPFC** |  |  |  |  |  |  |
| 10r | AntCing_MedPFC | L | 0.2887 | 0.8234* | [0.6068, 0.9262] | (-6.0, 60.0, -10.0) |
| 10v | AntCing_MedPFC | L | 0.4046 | 0.7647 | [0.4960, 0.8997] | (-2.0, 58.0, -14.0) |
| 25 | AntCing_MedPFC | L | 0.3789 | 0.6364 | [0.2809, 0.8383] | (0.0, 14.0, -10.0) |
| a24 | AntCing_MedPFC | L | 0.3500 | 0.6302 | [0.2713, 0.8352] | (-10.0, 40.0, -2.0) |
| a32pr | AntCing_MedPFC | L | 0.2325 | 0.5164 | [0.1076, 0.7759] | (-6.0, 32.0, 32.0) |
| d32 | AntCing_MedPFC | L | 0.2164 | 0.5427 | [0.1435, 0.7900] | (-2.0, 48.0, 28.0) |
| pOFC | AntCing_MedPFC | L | 0.2859 | 0.6294 | [0.2701, 0.8348] | (-22.0, 16.0, -20.0) |
| s32 | AntCing_MedPFC | L | 0.0506 | 0.3777 | [-0.0659, 0.6967] | (-6.0, 38.0, -18.0) |
| 11l | OrbPolaFrontal | L | 0.4066 | 0.8747* | [0.7111, 0.9485] | (-32.0, 46.0, -14.0) |
| 13l | OrbPolaFrontal | L | 0.3545 | 0.6611 | [0.3197, 0.8506] | (-16.0, 42.0, -18.0) |
| 47m | OrbPolaFrontal | L | 0.1669 | 0.5083 | [0.0967, 0.7714] | (-36.0, 38.0, -8.0) |
| OFC | OrbPolaFrontal | L | 0.3747 | 0.7401 | [0.4521, 0.8884] | (-10.0, 50.0, -20.0) |
| a47r | Inferior_Frontal | L | 0.2380 | 0.8152* | [0.5908, 0.9225] | (-34.0, 44.0, -14.0) |
| p47r | Inferior_Frontal | L | 0.4575 | 0.7957* | [0.5534, 0.9138] | (-50.0, 46.0, 6.0) |
| 10r | AntCing_MedPFC | R | 0.4498 | 0.7768* | [0.5182, 0.9053] | (8.0, 60.0, -6.0) |
| 10v | AntCing_MedPFC | R | 0.3477 | 0.6631 | [0.3229, 0.8515] | (6.0, 60.0, -10.0) |
| 25 | AntCing_MedPFC | R | 0.3437 | 0.7201 | [0.4174, 0.8790] | (8.0, 18.0, -14.0) |
| a24 | AntCing_MedPFC | R | 0.4474 | 0.7330 | [0.4397, 0.8851] | (8.0, 36.0, 0.0) |
| a32pr | AntCing_MedPFC | R | 0.4034 | 0.6062 | [0.2351, 0.8231] | (10.0, 22.0, 40.0) |
| d32 | AntCing_MedPFC | R | 0.2710 | 0.5345 | [0.1323, 0.7856] | (2.0, 46.0, 28.0) |
| pOFC | AntCing_MedPFC | R | 0.2187 | 0.7065 | [0.3941, 0.8725] | (10.0, 16.0, -16.0) |
| s32 | AntCing_MedPFC | R | 0.3177 | 0.6421 | [0.2898, 0.8412] | (6.0, 36.0, -20.0) |
| 11l | OrbPolaFrontal | R | 0.4876 | 0.8110* | [0.5827, 0.9207] | (32.0, 48.0, -14.0) |
| 13l | OrbPolaFrontal | R | 0.4388 | 0.7161 | [0.4104, 0.8771] | (26.0, 36.0, -10.0) |
| 47m | OrbPolaFrontal | R | 0.4668 | 0.7542 | [0.4771, 0.8949] | (30.0, 38.0, -6.0) |
| OFC | OrbPolaFrontal | R | 0.3975 | 0.7675* | [0.5011, 0.9010] | (12.0, 36.0, -20.0) |
| a47r | Inferior_Frontal | R | 0.3623 | 0.7508 | [0.4711, 0.8933] | (40.0, 62.0, -4.0) |
| p47r | Inferior_Frontal | R | 0.4551 | 0.8052* | [0.5715, 0.9181] | (44.0, 40.0, -4.0) |
| **Striatum** |  |  |  |  |  |  |
| Putam | Subcortical | L | 0.2666 | 0.8050* | [0.5711, 0.9180] | (-24.0, 18.0, -2.0) |
| Caud | Subcortical | L | 0.4187 | 0.8334* | [0.6266, 0.9306] | (-16.0, 18.0, 16.0) |
| NAc | Subcortical | L | 0.1478 | 0.4263 | [-0.0081, 0.7252] | (-12.0, 8.0, -10.0) |
| Putam | Subcortical | R | 0.3401 | 0.8310* | [0.6217, 0.9295] | (28.0, -12.0, 10.0) |
| Caud | Subcortical | R | 0.4933 | 0.8330* | [0.6258, 0.9304] | (18.0, 16.0, 4.0) |
| NAc | Subcortical | R | 0.3469 | 0.6318 | [0.2738, 0.8360] | (14.0, 8.0, -8.0) |

For parcels in each a priori ROI, the table summarizes the location of the most reliable voxel during feedback processing, together with its peak intraclass correlation coefficient (ICC) and MNI coordinates for BOLD responses in the delayed outcome condition. Asterisks indicate the reliability level of the peak ICC based on the lower bound of the 95% confidence interval: \* moderate reliability ( $> 0.50$ ), \*\* good reliability ( $> 0.75$ ), \*\*\* excellent reliability ( $> 0.90$ ).

For feedback-related activation, 7 of the 12 left DLPFC parcels had a 95% CI lower bound exceeding 0.50, indicating at least moderate reliability with confidence. Of these, one parcel, Area 8C, had a lower bound exceeding 0.75, confirming good reliability. The remaining five parcels, 8Ad, 8BL, 9-46d, 9a, and 9p, did not reach a lower bound of 0.50, indicating that their peak ICC estimates should be interpreted with caution.

For the extended regions during feedback-related activation, confirmed reliability differed across cortical divisions. In ACC/mPFC parcels, 1 of 8 left-hemisphere parcels and 1 of 8 right-hemisphere parcels exceeded a CI lower bound of 0.50. In OFC/vmPFC parcels, 3 of 6 left-hemisphere parcels and 3 of 6 right-hemisphere parcels exceeded this threshold. Striatal reliability was notably strong for caudate and putamen, with bilateral putamen and caudate each exceeding the 0.50 lower-bound threshold, corresponding to 4 of 6 total striatal parcels, while nucleus accumbens did not reach confirmed moderate reliability in either hemisphere.

## 4 Discussion

The main aim of this study was to assess the reliability of BOLD responses in the frontal-parietal and frontal-striatal networks during sequential decision making using a 3-stage Markov task. Behaviorally, we confirmed results of previous studies showing that a delayed temporal contingency between choices and outcomes across decision states resulted in lower performance and that initial performance level modulates the effect of temporal delays (Eppinger et al., 2015; Kang et al., 2024). In the whole-brain fMRI analysis, for the main effect of temporal delay (contrasting delayed with immediate condition), greater BOLD responses were observed in regions spanning the right insula and right frontal operculum, with additional involvement of striatal regions (e.g., bilateral putamen). Taken together, results of the reported task fMRI analyses are generally in line with previous studies showing the involvements of the frontal-parietal and frontal-striatal networks when learning choice-outcome associations during sequential decision making (Eppinger et al., 2015). Even in the relatively younger age range of this study (22 to 40 years), we observed a performance level by age interaction of activities in several frontal clusters (Figure 4), suggesting that high performers recruited greater activities in these regions when learning sequential action-outcome associations than lower performers, a finding that is in line with previous results assessed with functional near-infrared spectroscopy in older adults (Kang et al., 2024). The interaction with age indicates that this effect of performance modulation was mainly found in the older participants of this sample. Previous aging research showed that between-person differences in performance (Li et al., 2004) and activity patterns in task-relevant network tended to be larger in older ages (Nyberg et al., 2003). Taken together, these task-related functional results provide a basis for assessing the reliability of task-related BOLD responses.

Test-retest reliability analyses of task-related BOLD responses in the delayed condition (and for the delayed vs. immediate contrast, see supplementary materials) indicated numerous high voxel-wise point estimate ICC values (ranging from 0.34 to 0.89) in key regions. The reliability of these point estimates was strong (moderate to good) in multiple parcels in left DLPFC, and vmPFC/OFC parcels also showed multiple moderate to good peak ICC point estimates bilaterally. The ACC/mPFC parcels also exhibited some moderate-to-good point estimate reproducibility. Striatal regions (caudate, putamen, nucleus accumbens) showed strong and largely symmetric reliability across hemispheres, indicating consistent engagements of these reward-related structures in both scanning sessions.

### Frontal involvements in learning delayed action-outcome associations

We found higher BOLD responses in lateral DLPFC (including a left peak in the 9-46d/p47r region and a right peak in p9-46v/IFSp) in the delayed compared to the immediate condition. Previous studies of value-based sequential decision making using other variants of the three-stage Markov task (Eppinger et al., 2015; Tanaka et al., 2004) have shown that learning delayed action-outcome associations requires integrating temporally lagged action-outcome contingencies across multiple states, rather than relying on immediate outcome information. Converging evidence links left DLPFC to these processing demands, including acquiring abstract task rules and representing sequential relationships that support multi-step action planning (Badre & D’Esposito, 2007; Mansouri et al., 2020). Consistent with this interpretation, results from the exploratory High_Delay > Low_Immediate contrast also revealed bilateral DLPFC recruitment, particularly during the time period of feedback presentation, alongside additional dorsal frontal and fronto-cingulate peaks, suggesting that conditions combining delayed contingencies with high outcome magnitude further upregulate control-related circuitry.

Importantly, the present recruitment of left DLPFC is consistent with prior work using other variants of the Markov task, which also showed stronger prefrontal engagement when participants learn to predict future reward with delayed feedbacks (Tanaka et al., 2004), and with the evidence that learning state transitions across decision states in younger adults is initially accompanied by an increase BOLD responses in the left DLPFC. After the transition structure is learned, as indicated by performance change point, the activity of this region decreases (Eppinger et al., 2015). Therefore, the results involving the left DLPFC observed here are compatible with the interpretation that delayed outcomes increased reliance on adaptive control processes that maintain and update state information, enabling choices to be evaluated with respect to downstream consequences beyond outcomes of the current state.

We also observed greater activities around the vmPFC/bilateral OFC, alongside striatal involvement during the delayed relative to the immediate condition. Meta-analytic evidence suggests that vmPFC and ventral striatum are core components of a valuation system supporting subjective value representations (Bartra et al., 2013), and further research similarly highlights vmPFC-striatal circuitry for integrating value signals to guide choice (Rangel et al., 2008). In the context of the Markov task, these regions plausibly contribute to computing and updating policy values that depend on future outcomes, rather than on immediate reinforcement alone (Tanaka et al., 2004). A complementary account emphasizes OFC as representing task states or “cognitive maps” that disambiguate partially observable situations, which is especially relevant when early-stage outcomes are insufficient to determine long-term value and when optimal choice depends on anticipating later states (Wilson et al., 2014).

### Insular-opercular involvement as salience/control gating for delayed outcomes

Greater BOLD responses were observed in right insula and right frontal operculum during the delayed condition, with additional involvement of anterior ventral insular areas. This pattern is consistent with the engagement of the salience network, commonly anchored in anterior insula and anterior cingulate cortex (Menon & Uddin, 2010). One possible interpretation is that trials with delayed outcomes increase the need to detect and prioritize behaviorally relevant events under uncertainty, including the accumulation of small intermediate losses that must be tolerated to obtain a larger terminal gain. Within salience-network accounts, anterior insula is proposed to support bottom-up detection of salient events and to initiate switching between large-scale networks, including recruitment of frontoparietal control resources when demands increase (Menon & Uddin, 2010). This interpretation also aligns with previous studies of the Markov task. For instance, Tanaka et al. (2004) reported that insular and striatum differentiate predictions about immediate versus future rewards, with more anterior regions tracking immediate reward prediction and more posterior regions tracking future reward prediction. This account is also supported by exploratory analysis contrasting delayed high magnitude outcomes with immediate low magnitude outcomes, which again showed prominent insular–opercular engagement together with frontoparietal regions, consistent with increased salience and control allocation when delayed contingencies coincide with higher outcome magnitude

### Age moderates the performance-related recruitment of frontoparietal control circuits

In the whole-brain regressions, the Performance × Age term was positively associated with stimulus- and feedback-related BOLD responses in a distributed frontoparietal network. During stimulus processing, the interaction was most evident in bilateral DLPFC and additional orbitofrontal, parietal, and temporal regions, whereas during feedback it again involved DLPFC together with superior parietal regions, and in the Task2 extended to medial frontal areas. This pattern is broadly consistent with proposals suggesting that mid-DLPFC supports the integration of action–outcome contingencies across time in sequential decision settings (Badre & D’Esposito, 2007; Mansouri et al., 2020; Tanaka et al., 2004), and with prior evidence that age- or performance-related differences in Markov decision performance and intervention gains can be accompanied by differences in frontal recruitments (Eppinger et al., 2015; Kang et al., 2024).

### Measurement reliability of the Markov task

Accuracy and reaction time showed moderate to good test–retest reliability point estimates across the two task sessions, with ICC(3,1) values of 0.744 and 0.788, respectively. Interpreted within common reliability guidelines, these coefficients indicate that individual differences in performance were largely preserved between measurements, supporting the robustness of the behavioral indices in this sample. At the same time, ICC values are not solely properties of the task or measure but depend on the balance of between-subject variance relative to measurement error and other sources of within-subject variability. Because ICC(3,1) is a two-way mixed-effects consistency index, it reflects stable rank-ordering across sessions, implying that participants who showed stronger (or weaker) learning or more (or less) optimal choice tendencies in the first session tended to show comparatively stronger (or weaker) learning again in the second session, even if overall performance levels shifted between sessions (Shrout & Fleiss, 1979; Koo and Li, 2016).

Despite the overall decision accuracy showing a moderate ICC point estimate, its confidence interval suggests some uncertainty around the lower-bound reliability estimate, whereas overall mean reaction time showed more robust reliability with the lower confidence interval exceeding 0.50. This uncertainty likely reflects, at least in part, the individual differences in learning rate across task sessions and the modest sample size of the current study. However, together, generally these findings support the use of both behavioral indices in repeated-measures designs.

Establishing neural ICC reliability for BOLD responses during the Markov task under repeated sham tDCS provides an essential benchmark for the subsequent active stimulation phase of the overarching project. This is particularly relevant because task-based fMRI measures often show limited test–retest reliability, and low reliability directly constrains the detectability and interpretability of intervention effects (Caceres et al., 2009; Elliott et al., 2020; Noble et al., 2019). Within sequential decision making, converging evidence supports a functional role of the left DLPFC in integrating temporally extended action–outcome contingencies, and neuromodulation can alter left DLPFC hemodynamic responses during closely related Markov-task variants (Wittkuhn et al., 2018; Schommartz et al., 2021). Against this background, reliable sham–sham activation patterns in the target left DLPFC, alongside control stimulation regions such as the primary motor cortex in the follow-up stimulation groups, can help the active phase distinguish region-specific modulation from session-to-session variability driven by task design choices and participant factors such as motion or strategy variability (Bennett & Miller, 2010; Gorgolewski et al., 2012).

While point ICC estimates provide a useful summary of the central tendency of reliability, they can carry considerable uncertainty in small samples, and the 95% confidence intervals reported in this study offer a more conservative and informative basis for evaluating which regions can be deemed to reliably meet a given reliability threshold. Given the sample size of n = 20, confidence intervals were sometimes wide, and in several parcels the lower CI bound fell below 0.50 even where the peak ICC point estimate appeared moderate to good. This was particularly evident during stimulus presentation in the left DLPFC, where only 1 of 12 parcels had a lower bound exceeding 0.50. By contrast, during feedback processing, the majority (7 out of 12) of left DLPFC parcel peaks showed lower CI bounds consistently above 0.50, lending stronger inferential support to their reliability estimates. For intervention research, this distinction is consequential: regions whose reliability estimates are supported by the full confidence interval provide a firmer basis for detecting stimulation-induced modulation above the floor of session-to-session variability, whereas regions with wide or low-bounded CIs may require larger samples before their reliability can be confirmed with sufficient precision (Koo & Li, 2016; Elliott et al., 2020).

Importantly, confidence intervals quantify the precision of the ICC estimate but do not by themselves establish that a reliable signal is neural in origin. Thus, although the occasional high peak ICCs observed in certain parcels, in some cases exceeding point estimates of 0.8 and higher upper CI bounds upwards of 0.9, may reflect genuinely stable task-related BOLD responses across sessions, they could also partly reflect stable non-neural variance, including residual motion, and scanner-related noise (Bennett & Miller, 2010). We therefore view any of these peak point estimate values as encouraging but not definitive evidence of strong regional reliability.

### Conclusion

This study established that the deterministic three-stage Markov task can be implemented robustly in the current protocol and yields stable behavioral performance across repeated task sessions under sham stimulation. The observed test–retest consistency in accuracy and reaction time supports the use of these measures as baseline indices for the subsequent active stimulation phase, particularly when practice-related shifts are expected and relative performance differences are of primary interest (Shrout & Fleiss, 1979; Koo and Li, 2016). At the neural level, voxel-wise ICC mapping for the condition specific delayed regressor indicated many reproducible activation patterns in key circuits, especially during feedback presentation, previously linked to sequential value integration and control, providing a necessary benchmark for interpreting stimulation-related modulation against the background of session-to-session variability (Tanaka et al., 2004; Badre & D’Esposito, 2007; Rangel et al., 2008; Wittkuhn et al., 2018).

## Acknowledgments

This study took place in the initial preparatory phase of a project funded by the German Research Foundation (DFG; project grants: Research Unit 5429/1, [467143400]; Subproject P8 LI879/24-1). We thank Dr. Till Nierhaus for his technical support in preparing the imaging and stimulation setup. We also thank Oona Lufft for her assistance with data collection.

## Ethics Statement

This research was conducted at the Center for Cognitive Neuroscience Berlin (CCNB) and approved by the Ethics Committee of Technische Universität Dresden (SR-EK-229052022). The Ethics Committee of the Department of Education and Psychology at the Freie Universität Berlin also accepted the ethical approval given by TU Dresden (No. 045/2022) and for using the imaging facility at the CCNB.

## Author Contributions

NLS: Methodology, Software, Validation, Formal analysis, Investigation, Data Curation, Writing – Original Draft, Writing – Review & Editing, Visualization. JP: Methodology, Software, Validation, Formal analysis, Data curation, Writing – Review & Editing, Visualization. MJ: Investigation, Data curation, Project administration. FN: Software (initial codes for ICC analyses). MG: Formal analysis (consultation). IMG: Visualization. YS: Conceptualization, Methodology, Software, Investigation, Supervision. SCL: Conceptualization, Methodology, Resources, Supervision, Writing – Review & Editing, Funding acquisition.

## Data Availability Statement

This study was pre-registered on the Open Science Framework (OSF). The pre-registration materials can be accessed under OSF Registries (https://osf.io/t37u2). The data reported in this study are available upon request from researchers by contacting the corresponding authors. Due to personal data privacy restrictions in ethical approval, the data is not publicly available in an open-source form.

