## Supplementary Materials for "Retest Reliability of Task-related fMRI BOLD Signals during Sequential Decision Making"

In total three complementary TRR analyses of task-related BOLD responses are reported in the supplementary materials. The first analysis used condition-specific beta estimates for the delayed condition, identical to the one in the main manuscript however, omitting the minimum voxel coverage criterion and using PyReliMRI's default handling of missing observations.

Furthermore, we also assessed TRR using ICC computed based on beta values of the main effect contrast (delayed > immediate) from the first-level analysis and assessed TRR without the minimum voxel criterion.

For each a priori parcel of interest previously implicated in value-based learning, the peak ICC voxel and its MNI coordinates, the 95% confidence interval at that peak, and the parcel-level mean ICC are reported. Of these, one retained only voxels with valid data from at least 10 participants, whereas the other applied no minimum voxel coverage criterion and handled missing observations using PyReliMRIs (Demidenko et al., 2024) default column-mean imputation procedure.

TRR Regressor Specific Results (PyReliMRI's default method) - Stimulus Presentation

As part of the condition-specific analysis with no minimum voxel coverage criterion, TRR of BOLD responses during stimulus presentation was assessed by computing ICC maps for the delayed condition-specific regressor and overlaying them with the HCPex atlas.

Left dorsolateral prefrontal cortex (DLPFC) parcels demonstrated moderate-to-good peak ICC point estimates. The highest reliability was found in Area 46 (MNI: -38, 48, 32; ICC = 0.849) and Area 8BL (MNI: -10, 38, 60; ICC = 0.793), followed by Area 9a (MNI: -28, 56, 32; ICC = 0.757) and Area 8C (MNI: -40, 0, 38; ICC = 0.721).

ICC values for all key regions are presented in Table S1 and summarized as an overview in Figure S1, left panel.

In the vmPFC/OFC parcels, peak ICC point estimates ranged from 0.513 to 0.999 and were predominantly high, with excellent values observed in the left OFC, left 11l, and right OFC. ACC/mPFC parcels also showed some excellent point estimate peaks  such as left Area 10v (MNI: 0, 52, -24; ICC = 0.997) and right 10v (MNI: 8, 68, -14; ICC = 0.999), left pOFC (MNI: -4, 12, -22; ICC = 0.999), right pOFC (MNI: 10, 16, -20; ICC = 0.992) and overall point estimates ranged from 0.341 to 0.999.

Based on ICC point estimates, striatal parcels showed generally strong and symmetric reliability across hemispheres. In the left hemisphere, putamen (MNI: -28, 8, 8; ICC = 0.752) and caudate (MNI: -8, 10, 16; ICC = 0.761) demonstrated good peak reliability, whereas nucleus accumbens showed poor reliability (MNI: -6, 8, -12; ICC = 0.483). In the right hemisphere, reliability was similarly high for caudate (MNI: 14, 6, 22; ICC = 0.825) and putamen (MNI: 28, 16, -4; ICC = 0.779), whereas nucleus accumbens showed moderate reliability (MNI: 14, 8, -8; ICC = 0.601).

For stimulus-related activation in the delayed condition, confidence interval-based evaluation indicated that confirmed reliability was concentrated in a subset of parcels. Within left DLPFC, 2 of 12 parcels had a 95% confidence interval lower bound exceeding 0.50: Area 46 and Area 8BL. Among the extended cortical regions, confirmed reliability differed across cortical divisions. In ACC/mPFC parcels, 3 of 8 left-hemisphere parcels and 5 of 8 right-hemisphere parcels exceeded a lower confidence interval bound of 0.50. In vmPFC/OFC parcels, 3 of 6 parcels per hemisphere exceeded this threshold. Striatal parcels showed limited confirmed reliability, with 2 of 6 parcels exceeding the 0.50 lower-bound threshold, specifically right putamen and right caudate.

TRR Regressor Specific Results  (PyReliMRI's default method) - Feedback Presentation

As part of the condition-specific analysis with no minimum voxel coverage criterion, TRR of BOLD responses during feedback presentation was assessed by computing ICC maps for the delayed condition-specific regressor and overlaying them with the HCPex atlas.

Left DLPFC parcels showed predominantly high peak ICC point estimates. Excellent reliability was observed in Area 9a (MNI: -16, 68, 22; ICC = 0.950), whereas Area 8C (MNI: -40, 4, 34; ICC = 0.899) and Area 46 (MNI: -42, 46, 30; ICC =

0.897) demonstrated good reliability. The lowest point estimate reliability within the examined left DLPFC parcels was observed in Area 8Ad (MNI: -24, 28, 40; ICC = 0.609), which nevertheless indicated moderate reliability. ICC values for all key regions are presented in Table S2 and summarized in Figure S1, right panel.

Based on ICC point estimates, striatal parcels showed strong and symmetric reliability across hemispheres. In the left hemisphere, caudate (MNI: -16, 18, 16; ICC = 0.833) and putamen (MNI: -24, 18, -2; ICC = 0.804) demonstrated good peak reliability, whereas nucleus accumbens showed moderate reliability (MNI: -8, 10, -12; ICC = 0.703). In the right hemisphere, comparable reliability was observed in caudate (MNI: 18, 16, 4; ICC = 0.833) and putamen (MNI: 28, -12, 10; ICC = 0.830), whereas nucleus accumbens showed moderate reliability (MNI: 14, 8, -8; ICC = 0.631).

For feedback-related activation in the delayed condition, confidence interval-based evaluation showed stronger confirmed reliability in left DLPFC than during stimulus presentation. Within left DLPFC, 9 of 12 parcels had a 95% confidence interval lower bound exceeding 0.50, indicating at least moderate reliability. Of these, three parcels had lower bounds exceeding 0.75: Area 46 (lower bound = 0.759), Area 8C (lower bound = 0.763), and Area 9a (lower bound = 0.880), supporting good-to-excellent reliability.

Among the extended cortical regions, confirmed reliability differed across cortical divisions during feedback presentation. In ACC/mPFC parcels, 4 of 8 left-hemisphere parcels and 4 of 8 right-hemisphere parcels exceeded a lower confidence interval bound of 0.50. In OFC/vmPFC parcels, 5 of 6 left-hemisphere parcels and 3 of 6 right-hemisphere parcels exceeded this threshold. Striatal reliability was notably strong, with bilateral putamen and caudate exceeding the 0.50 lower-bound threshold, corresponding to 4 of 6 striatal parcels, whereas nucleus accumbens did not reach confirmed moderate reliability in either hemisphere.

***Figure S1. Whole-brain voxel-wise heat map of TRR across task sessions for BOLD responses of the condition specific regressor (delayed).***

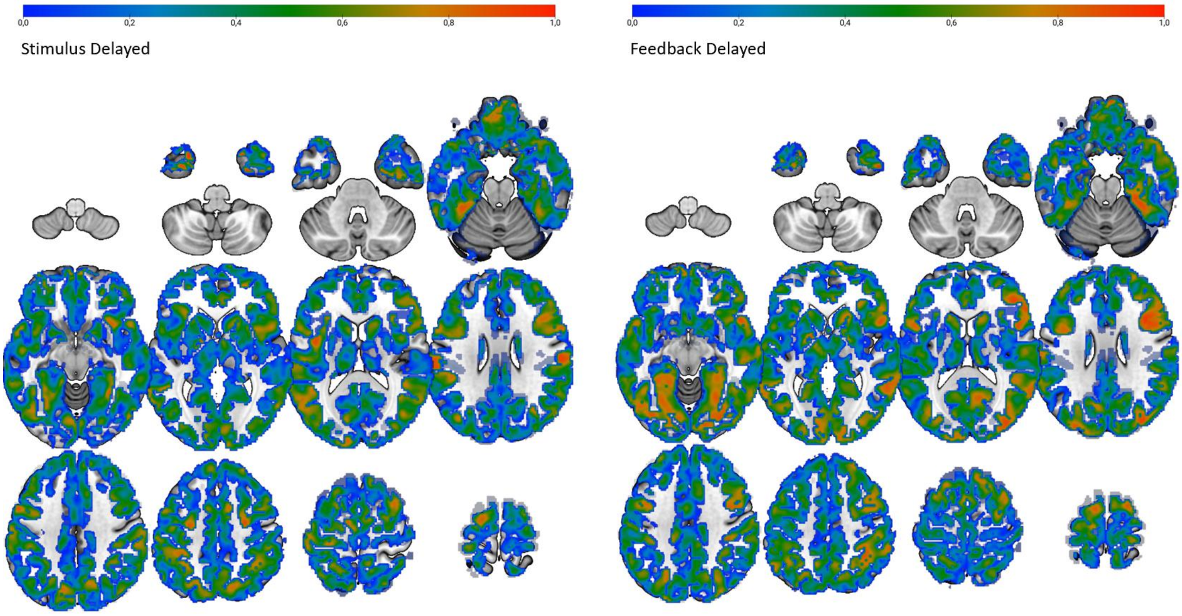


*Whole-brain voxel-wise ICC map computed for the delayed condition, for stimulus-related activity (left) and feedback-related activity (right). Only regions that fall under the HCPex atlas are shown.*

***Table S1. Peak ICC values of key region defined by the HCPex atlas during stimulus presentation for the regressor specific beta (delayed condition) using PyReliMRI's default method.***

| Parcel | Cortical Division | Side | Mean ICC | Peak ICC | 95% CI | Peak MNI (x, y, z) |
|---|---|---|---|---|---|---|
| | | | DLPFC | | | |
| 46 | Dorsolateral_Prefrontal | L | 0.3463 | 0.8496* | [0.6590, 0.9380] | (-38.0, 48.0, 32.0) |
| 8Ad | Dorsolateral_Prefrontal | L | 0.0801 | 0.6001 | [0.2260, 0.8200] | (-32.0, 32.0, 52.0) |
| 8Av | Dorsolateral_Prefrontal | L | 0.2422 | 0.5682 | [0.1800, 0.8030] | (-44.0, 26.0, 42.0) |
| 8BL | Dorsolateral_Prefrontal | L | 0.3756 | 0.7937* | [0.5500, 0.9130] | (-10.0, 38.0, 60.0) |
| 8C | Dorsolateral_Prefrontal | L | 0.4880 | 0.7217 | [0.4200, 0.8800] | (-40.0, 0.0, 38.0) |
| 9-46d | Dorsolateral_Prefrontal | L | 0.1807 | 0.6659 | [0.3270, 0.8530] | (-34.0, 52.0, 32.0) |
| 9a | Dorsolateral_Prefrontal | L | 0.1233 | 0.7578 | [0.4840, 0.8970] | (-28.0, 56.0, 32.0) |
| 9p | Dorsolateral_Prefrontal | L | 0.3095 | 0.6640 | [0.3240, 0.8520] | (-12.0, 52.0, 30.0) |
| a9-46v | Dorsolateral_Prefrontal | L | 0.3050 | 0.6236 | [0.2610, 0.8320] | (-42.0, 50.0, 22.0) |
| i6-8 | Dorsolateral_Prefrontal | L | 0.3466 | 0.5822 | [0.2000, 0.8110] | (-24.0, 6.0, 60.0) |
| s6-8 | Dorsolateral_Prefrontal | L | 0.3289 | 0.6761 | [0.3439, 0.8580] | (-22.0, 22.0, 62.0) |
| SFL | Dorsolateral_Prefrontal | L | 0.2324 | 0.6280 | [0.2680, 0.8340] | (-8.0, 2.0, 70.0) |
| | | | vmPFC/OFC and ACC/mPFC | | | |
| 10r | AntCing_MedPFC | L | 0.3518 | 0.9185** | [0.8063, 0.9669] | (-2.0, 62.0, -6.0) |
| 10v | AntCing_MedPFC | L | 0.4504 | 0.9973*** | [0.9932, 0.9989] | (0.0, 52.0, -24.0) |
| 25 | AntCing_MedPFC | L | 0.2670 | 0.7460 | [0.4625, 0.8911] | (-4.0, 14.0, -16.0) |
| a24 | AntCing_MedPFC | L | 0.4340 | 0.6628 | [0.3225, 0.8514] | (0.0, 44.0, -6.0) |
| a32pr | AntCing_MedPFC | L | 0.1964 | 0.3414 | [-0.1072, 0.6746] | (-8.0, 24.0, 36.0) |
| d32 | AntCing_MedPFC | L | 0.2275 | 0.5152 | [0.1060, 0.7752] | (-6.0, 50.0, 32.0) |
| pOFC | AntCing_MedPFC | L | 0.1153 | 0.9998*** | [0.9996, 0.9999] | (-4.0, 12.0, -22.0) |
| s32 | AntCing_MedPFC | L | 0.1395 | 0.4361 | [0.0040, 0.7310] | (-8.0, 40.0, -8.0) |
| 11l | OrbPolaFrontal | L | 0.3885 | 0.9996*** | [0.9991, 0.9998] | (-20.0, 58.0, -18.0) |
| 13l | OrbPolaFrontal | L | 0.2577 | 0.7619 | [0.4909, 0.8984] | (-16.0, 44.0, -20.0) |
| 47m | OrbPolaFrontal | L | 0.2330 | 0.5134 | [0.1035, 0.7742] | (-28.0, 36.0, -14.0) |
| OFC | OrbPolaFrontal | L | 0.3538 | 0.9998*** | [0.9997, 0.9999] | (-6.0, 36.0, -26.0) |
| a47r | Inferior_Frontal | L | 0.2603 | 0.9046** | [0.7753, 0.9611] | (-38.0, 64.0, 0.0) |
| p47r | Inferior_Frontal | L | 0.2785 | 0.5867 | [0.2063, 0.8131] | (-46.0, 40.0, 6.0) |
| 10r | AntCing_MedPFC | R | 0.4461 | 0.8156* | [0.5916, 0.9227] | (6.0, 62.0, -6.0) |
| 10v | AntCing_MedPFC | R | 0.4400 | 0.9996*** | [0.9990, 0.9998] | (8.0, 68.0, -14.0) |
| 25 | AntCing_MedPFC | R | 0.2006 | 0.8538* | [0.6677, 0.9394] | (2.0, 10.0, -14.0) |
| a24 | AntCing_MedPFC | R | 0.5255 | 0.7693* | [0.5045, 0.9018] | (6.0, 38.0, -2.0) |
| a32pr | AntCing_MedPFC | R | 0.3192 | 0.5705 | [-0.1828, 0.8046] | (10.0, 22.0, 40.0) |
| d32 | AntCing_MedPFC | R | 0.1848 | 0.4325 | [-0.0003, 0.7289] | (6.0, 40.0, 34.0) |
| pOFC | AntCing_MedPFC | R | 0.1355 | 0.9922*** | [0.9805, 0.9969] | (10.0, 16.0, -20.0) |
| s32 | AntCing_MedPFC | R | 0.2936 | 0.5427 | [0.1435, 0.7899] | (2.0, 42.0, -10.0) |
| 11l | OrbPolaFrontal | R | 0.3426 | 0.8397* | [0.6392, 0.9333] | (18.0, 40.0, -20.0) |
| 13l | OrbPolaFrontal | R | 0.3255 | 0.7257 | [0.4269, 0.8816] | (20.0, 42.0, -18.0) |
| 47m | OrbPolaFrontal | R | 0.4249 | 0.7184 | [0.4145, 0.8782] | (30.0, 38.0, -6.0) |
| OFC | OrbPolaFrontal | R | 0.2982 | 0.9998*** | [0.9997, 0.9999] | (10.0, 54.0, -26.0) |
| a47r | Inferior_Frontal | R | 0.2534 | 0.7729* | [0.5110, 0.9034] | (28.0, 52.0, -4.0) |
| p47r | Inferior_Frontal | R | 0.4474 | 0.6868 | [0.3614, 0.8630] | (46.0, 40.0, -6.0) |
| | | | Striatum | | | |
| Putam | Subcortical | L | 0.2997 | 0.7524 | [0.4739, 0.8940] | (-28.0, 8.0, 8.0) |
| Caud | Subcortical | L | 0.3395 | 0.7613 | [0.4900, 0.8982] | (-8.0, 10.0, 16.0) |
| NAc | Subcortical | L | -0.2091 | 0.4834 | [0.0639, 0.7577] | (-6.0, 8.0, -12.0) |
| Putam | Subcortical | R | 0.3692 | 0.7795* | [0.5232, 0.9065] | (28.0, 16.0, -4.0) |
| Caud | Subcortical | R | 0.4296 | 0.8258* | [0.6115, 0.9272] | (14.0, 6.0, 22.0) |
| NAc | Subcortical | R | -0.0033 | 0.6019 | [0.2286, 0.8209] | (14.0, 8.0, -8.0) |

*For each a priori region of interest, the table summarizes the location of the most reliable voxel during stimulus processing, together with its peak intraclass correlation coefficient (ICC) and MNI coordinates for the delayed specific regressor. Asterisks indicate the reliability level of the peak ICC based on the lower bound of the 95% confidence interval: * moderate reliability (> 0.50), ** good reliability (> 0.75), *** excellent reliability (> 0.90).*

***Table S2. Peak ICC values of key region defined by the HCPex atlas during feedback presentation for the regressor specific beta (delayed condition) using PyReliMRI's default method.***

| Parcel | Cortical Division | Side | Mean ICC | Peak ICC | 95% CI | Peak MNI (x, y, z) |
|---|---|---|---|---|---|---|
| | | | DLPFC | | | |
| 46 | Dorsolateral_Prefrontal | L | 0.3145 | 0.8972** | [0.7590, 0.9580] | (-42.0, 46.0, 30.0) |
| 8Ad | Dorsolateral_Prefrontal | L | 0.3305 | 0.6095 | [0.2400, 0.8250] | (-24.0, 28.0, 40.0) |
| 8Av | Dorsolateral_Prefrontal | L | 0.3020 | 0.7910* | [0.5450, 0.9120] | (-30.0, 14.0, 60.0) |
| 8BL | Dorsolateral_Prefrontal | L | 0.3759 | 0.7403 | [0.4530, 0.8890] | (-8.0, 38.0, 56.0) |
| 8C | Dorsolateral_Prefrontal | L | 0.3483 | 0.8990** | [0.7630, 0.9590] | (-40.0, 4.0, 34.0) |
| 9-46d | Dorsolateral_Prefrontal | L | 0.3691 | 0.7943* | [0.5510, 0.9130] | (-40.0, 52.0, 24.0) |
| 9a | Dorsolateral_Prefrontal | L | 0.2200 | 0.9509** | [0.8800, 0.9800] | (-16.0, 68.0, 22.0) |
| 9p | Dorsolateral_Prefrontal | L | 0.3585 | 0.7663 | [0.4990, 0.9000] | (-28.0, 50.0, 38.0) |
| a9-46v | Dorsolateral_Prefrontal | L | 0.2481 | 0.8062* | [0.5729, 0.9180] | (-44.0, 48.0, 22.0) |
| i6-8 | Dorsolateral_Prefrontal | L | 0.4531 | 0.8001* | [0.5620, 0.9160] | (-30.0, 12.0, 60.0) |
| s6-8 | Dorsolateral_Prefrontal | L | 0.4872 | 0.7808* | [0.5260, 0.9070] | (-20.0, 24.0, 66.0) |
| SFL | Dorsolateral_Prefrontal | L | 0.5254 | 0.8176* | [0.5950, 0.9240] | (-10.0, 16.0, 72.0) |
| | | | vmPFC/OFC and ACC/mPFC | | | |
| 10r | AntCing_MedPFC | L | 0.3048 | 0.9544** | [0.8888, 0.9817] | (-4.0, 62.0, -6.0) |
| 10v | AntCing_MedPFC | L | 0.4033 | 0.9994*** | [0.9986, 0.9997] | (-4.0, 38.0, -26.0) |
| 25 | AntCing_MedPFC | L | 0.3850 | 0.8446* | [0.6491, 0.9354] | (-4.0, 14.0, -16.0) |
| a24 | AntCing_MedPFC | L | 0.3499 | 0.6301 | [0.2713, 0.8352] | (-10.0, 40.0, -2.0) |
| a32pr | AntCing_MedPFC | L | 0.2325 | 0.5164 | [0.1076, 0.7758] | (-6.0, 32.0, 32.0) |
| d32 | AntCing_MedPFC | L | 0.2163 | 0.5426 | [0.1435, 0.7899] | (-2.0, 48.0, 28.0) |
| pOFC | AntCing_MedPFC | L | 0.0739 | 0.9992*** | [0.9954, 0.9992] | (-14.0, 12.0, -22.0) |
| s32 | AntCing_MedPFC | L | 0.0506 | 0.3777 | [-0.0659, 0.6966] | (-6.0, 38.0, -18.0) |
| 11l | OrbPolaFrontal | L | 0.3923 | 0.9516** | [0.8822, 0.9805] | (-20.0, 58.0, -18.0) |
| 13l | OrbPolaFrontal | L | 0.3052 | 0.7676* | [0.5014, 0.9011] | (-24.0, 20.0, -24.0) |
| 47m | OrbPolaFrontal | L | 0.1669 | 0.5082 | [0.0967, 0.7714] | (-36.0, 38.0, -8.0) |
| OFC | OrbPolaFrontal | L | 0.3674 | 0.9999*** | [0.9999, 0.9999] | (-4.0, 34.0, -26.0) |
| a47r | Inferior_Frontal | L | 0.2430 | 0.8220* | [0.6042, 0.9255] | (-36.0, 62.0, -10.0) |
| p47r | Inferior_Frontal | L | 0.4568 | 0.7956* | [0.5533, 0.9137] | (-50.0, 46.0, 6.0) |
| 10r | AntCing_MedPFC | R | 0.4701 | 0.8308* | [0.6215, 0.9294] | (6.0, 62.0, -6.0) |
| 10v | AntCing_MedPFC | R | 0.4085 | 0.9877*** | [0.9694, 0.9951] | (8.0, 68.0, -14.0) |
| 25 | AntCing_MedPFC | R | 0.3477 | 0.8247* | [0.6095, 0.9267] | (2.0, 14.0, -20.0) |
| a24 | AntCing_MedPFC | R | 0.4474 | 0.7330 | [0.4396, 0.8850] | (8.0, 36.0, 0.0) |
| a32pr | AntCing_MedPFC | R | 0.4033 | 0.6062 | [0.2350, 0.8231] | (10.0, 22.0, 40.0) |
| d32 | AntCing_MedPFC | R | 0.2709 | 0.5345 | [0.1322, 0.7856] | (2.0, 46.0, 28.0) |
| pOFC | AntCing_MedPFC | R | 0.2694 | 0.9999*** | [0.9997, 0.9999] | (14.0, 14.0, -18.0) |
| s32 | AntCing_MedPFC | R | 0.3177 | 0.6420 | [0.2897, 0.8411] | (6.0, 36.0, -20.0) |
| 11l | OrbPolaFrontal | R | 0.4170 | 0.8109* | [0.5826, 0.9206] | (32.0, 48.0, -14.0) |
| 13l | OrbPolaFrontal | R | 0.3839 | 0.7287 | [0.4321, 0.8830] | (20.0, 30.0, -24.0) |
| 47m | OrbPolaFrontal | R | 0.4667 | 0.7542 | [0.4771, 0.8949] | (30.0, 38.0, -6.0) |
| OFC | OrbPolaFrontal | R | 0.3774 | 0.9998*** | [0.9995, 0.9999] | (2.0, 52.0, -24.0) |
| a47r | Inferior_Frontal | R | 0.3642 | 0.7508 | [0.4710, 0.8933] | (40.0, 62.0, -4.0) |
| p47r | Inferior_Frontal | R | 0.4550 | 0.8051* | [0.5714, 0.9180] | (44.0, 40.0, -4.0) |
| | | | Striatum | | | |
| Putam | Subcortical | L | 0.2694 | 0.8049* | [0.5711, 0.9179] | (-24.0, 18.0, -2.0) |
| Caud | Subcortical | L | 0.4187 | 0.8334* | [0.6266, 0.9305] | (-16.0, 18.0, 16.0) |
| NAc | Subcortical | L | 0.1565 | 0.7039 | [0.3899, 0.8713] | (-8.0, 10.0, -12.0) |
| Putam | Subcortical | R | 0.3401 | 0.8309* | [0.6217, 0.9294] | (28.0, -12.0, 10.0) |
| Caud | Subcortical | R | 0.4933 | 0.8330* | [0.6257, 0.9303] | (18.0, 16.0, 4.0) |
| NAc | Subcortical | R | 0.3166 | 0.6317 | [0.2737, 0.8360] | (14.0, 8.0, -8.0) |

*For each a priori region of interest, the table summarizes the location of the most reliable voxel during feedback processing, together with its peak intraclass correlation coefficient (ICC) and MNI coordinates for the delayed specific regressor. Asterisks indicate the reliability level of the peak ICC based on the lower bound of the 95% confidence interval: * moderate reliability (> 0.50), ** good reliability (> 0.75), *** excellent reliability (> 0.90).*

<u>TRR Contrast Results (PyReliMRI's default method) - Stimulus Presentation</u>

As part of the main effect contrast analysis with no minimum voxel coverage criterion (missing observations handled via PyReliMRI's default column-mean imputation), TRR of BOLD responses during stimulus presentation was assessed by computing ICC maps for the delayed > immediate contrast and overlaying them with the HCPex atlas.

Left DLPFC parcels demonstrated weak-to-excellent peak ICC points estimates overall, ranging from 0.430 to 0.964. The highest reliability was found in area 8Av (MNI: -32, 22, 60; ICC = 0.964), followed by area 46 (MNI: -32, 40, 46; ICC = 0.845), area 8C (MNI: -52, 14, 46; ICC = 0.837), and area 8BL (MNI: -22, 30, 60; ICC = 0.808).

In the vmPFC/OFC parcels, peak ICC point estimates ranged from 0.2124 to 0.999 and were predominantly high, with excellent values observed in left OFC, right 11l, right 13l, and right OFC. ACC/mPFC parcels showed excellent point estimates in parcel 10v and parcel pOFC of both hemispheres. ICC values for all key regions are presented in Table S3 and summarized as an overview in Figure S2 (left panel).

Striatal parcels showed weak-to-moderate ICC point estimates across hemispheres, with peak ICCs ranging from 0.374 to 0.735. For instance, on the left, nucleus accumbens (MNI: -8, 8, -12; ICC = 0.619) and putamen (MNI: -16, 10, -14; ICC = 0.559) showed fair peak reliability. On the right, reliability was highest for caudate (MNI: 26, -32, 12; ICC = 0.735), followed by nucleus accumbens (MNI: 10, 10, -12; ICC = 0.605 ), whereas putamen showed weak reliability (MNI: 24, 6, 14; ICC = 0.374).

For the stimulus-related delayed > immediate contrast, confidence interval-based evaluation indicated that confirmed reliability was concentrated in a subset of parcels. Within left DLPFC, 4 of 12 parcels had a 95% CI lower bound exceeding 0.50: Area 46, 8Av, 8BL, and 8C. Of these, Area 8Av had a lower bound exceeding 0.90, supporting excellent reliability. Among the extended cortical regions, confirmed reliability differed across cortical divisions.

In ACC/mPFC parcels, 2 of 8 left-hemisphere parcels and 2 of 8 right-hemisphere parcels exceeded a lower CI bound of 0.50. In vmPFC/OFC parcels, 2 of 6 left-hemisphere parcels and 3 of 6 right-hemisphere parcels exceeded this threshold. No striatal parcel exceeded a lower CI bound of 0.50.

TRR Contrast Results (PyReliMRI’s default method) - Feedback Presentation

As part of the main effect contrast analysis with no minimum voxel coverage criterion (missing observations handled via PyReliMRI's default column-mean imputation), TRR of BOLD responses during feedback presentation was assessed by computing ICC maps for the delayed > immediate contrast and overlaying them with the HCPex atlas.

DLPFC parcels showed weak-to-excellent peak ICCs overall, ranging from 0.213 to 0.991. The highest reliability was observed in Area 8Av (MNI: -40, 26, 52; ICC = 0.991), followed by Area 8BL (MNI: -8, 42, 50; ICC = 0.535), Area 9a (MNI: -26, 54, 12; ICC = 0.529), and Area 9-46d (MNI: -26, 50, 6; ICC = 0.524).

In the vmPFC/OFC parcels, peak ICC point estimates ranged from 0.199 to 0.999 and were particularly high in bilateral OFC regions. The strongest reliabilities were observed in left OFC (MNI: -4, 34, -26; ICC = 0.999), right OFC (MNI: 16, 30, -24; ICC = 0.999), and left 11l (MNI: -28, 66, -14; ICC = 0.982). By contrast, ACC/mPFC parcels also showed peak ICC point estimates of 0.999. The highest values in this region were observed in left pOFC (MNI: -4, 12, -20; ICC = 0.999), right pOFC (MNI: 14, 14, -20; ICC = 0.986), right Area 25 (MNI: 2, 14, -20; ICC = 0.718), right 10v (MNI: 4, 66, -16; ICC = 0.967), and left 10v (MNI: -6, 66, -8; ICC = 0.986). ICC values for all key regions are presented in Table S4 and summarized in Figure S2 (right panel).

Striatal parcels showed weak-to-moderate peak reliability, with ICC point estimates ranging from -0.036 to 0.639. In the left hemisphere, caudate showed the highest striatal reliability (MNI: -16, -12, 26; ICC = 0.580), followed by putamen (MNI: -28, 2, 0; ICC = 0.534), whereas nucleus accumbens showed negative reliability (MNI: -16, 8, -14; ICC = -0.036). In the right hemisphere, reliability was highest for caudate (MNI: 26, -32, 12; ICC = 0.639), followed by putamen (MNI: 22, 20, 2; ICC = 0.571) and nucleus accumbens (MNI: 10, 8, -4; ICC = 0.181).

For the feedback-related delayed > immediate contrast, confidence interval-based evaluation showed confirmed reliability primarily in selected ACC/mPFC and vmPFC/OFC parcels. Within left DLPFC, only 1 of 12 parcels, Area 8Av, had a 95% CI lower bound exceeding 0.50, and this lower bound also exceeded 0.90, supporting excellent reliability. Among the extended cortical regions, confirmed reliability differed across cortical divisions. In ACC/mPFC parcels, 2 of 8 left-hemisphere parcels and 3 of 8 right-hemisphere parcels exceeded a lower CI bound of 0.50. In vmPFC/OFC parcels, 2 of 6 left-hemisphere parcels and 3 of 6 right-hemisphere parcels exceeded this threshold. No striatal parcel exceeded a lower CI bound of 0.50.

***Figure S2. Whole-brain voxel-wise heat map of TRR across task sessions for BOLD responses of the main effect contrast (delayed > immediate)***

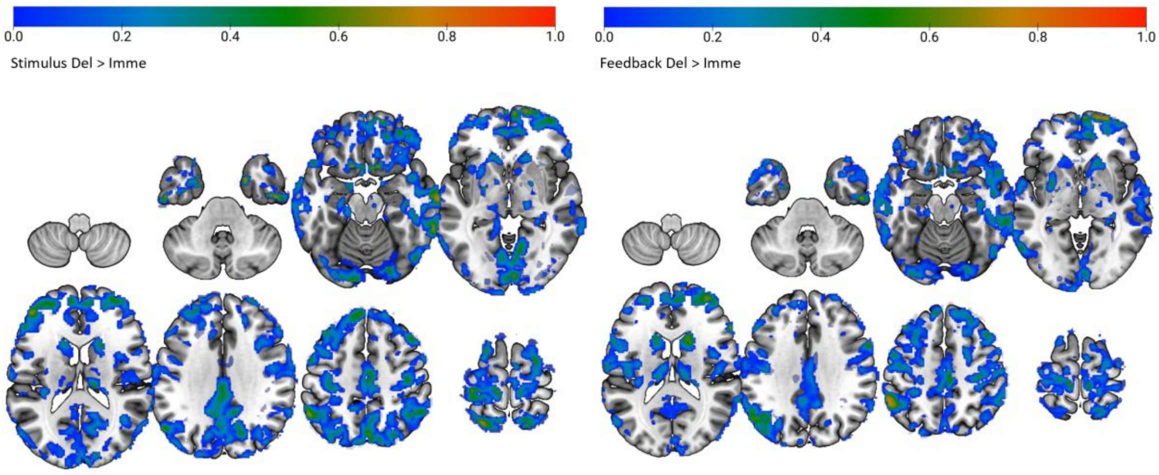


*Whole-brain voxel-wise ICC map computed for Delayed > Immediate for stimulus-related activity (left) and feedback-related activity (right) across the two task sessions.*

***Table S3. Peak ICC values of key region defined by the HCPex atlas during stimulus presentation for the main effect contrast (delayed > immediate) using PyReliMRI's default method.***

| Parcel | Cortical Division | Side | Mean ICC | Peak ICC | 95% CI | Peak MNI (x, y, z) |
|---|---|---|---|---|---|---|
| | | | DLPFC | | | |
| 46 | Dorsolateral_Prefrontal | L | 0.0237 | 0.8458* | [0.6510, 0.9360] | (-32.0, 40.0, 46.0) |
| 8Ad | Dorsolateral_Prefrontal | L | 0.1583 | 0.5905 | [0.2120, 0.8149] | (-32.0, 32.0, 52.0) |
| 8Av | Dorsolateral_Prefrontal | L | 0.2319 | 0.9643*** | [0.9120, 0.9860] | (-32.0, 22.0, 60.0) |
| 8BL | Dorsolateral_Prefrontal | L | 0.1842 | 0.8080* | [0.5770, 0.9190] | (-22.0, 30.0, 60.0) |
| 8C | Dorsolateral_Prefrontal | L | -0.0983 | 0.8375* | [0.6350, 0.9320] | (-52.0, 14.0, 46.0) |
| 9-46d | Dorsolateral_Prefrontal | L | 0.0651 | 0.5750 | [0.1890, 0.8070] | (-24.0, 54.0, 12.0) |
| 9a | Dorsolateral_Prefrontal | L | 0.0777 | 0.7259 | [0.4270, 0.8820] | (-20.0, 60.0, 32.0) |
| 9p | Dorsolateral_Prefrontal | L | 0.0639 | 0.4636 | [0.0380, 0.7470] | (-24.0, 48.0, 34.0) |
| a9-46v | Dorsolateral_Prefrontal | L | 0.2632 | 0.7255 | [0.4270, 0.8820] | (-48.0, 46.0, 14.0) |
| i6-8 | Dorsolateral_Prefrontal | L | -0.0106 | 0.4305 | [-0.0030, 0.7280] | (-40.0, 2.0, 62.0) |
| s6-8 | Dorsolateral_Prefrontal | L | 0.1501 | 0.6893 | [0.3650, 0.8640] | (-26.0, 20.0, 64.0) |
| SFL | Dorsolateral_Prefrontal | L | -0.3277 | 0.4709 | [0.0480, 0.7510] | (-8.0, 24.0, 68.0) |
| | | | vmPFC/OFC and ACC/mPFC | | | |
| 10r | AntCing_MedPFC | L | 0.0210 | 0.5046 | [0.0918, 0.7694] | (-10.0, 48.0, -8.0) |
| 10v | AntCing_MedPFC | L | 0.0195 | 0.9740*** | [0.9356, 0.9896] | (-6.0, 66.0, -10.0) |
| 25 | AntCing_MedPFC | L | -0.1504 | 0.4182 | [-0.0178, 0.7206] | (-4.0, 14.0, -16.0) |
| a24 | AntCing_MedPFC | L | 0.0487 | 0.5093 | [0.0980, 0.7719] | (-2.0, 50.0, -6.0) |
| a32pr | AntCing_MedPFC | L | -0.1092 | 0.2494 | [-0.2056, 0.6157] | (-10.0, 26.0, 34.0) |
| d32 | AntCing_MedPFC | L | -0.1082 | 0.3616 | [-0.0843, 0.6869] | (-14.0, 48.0, 10.0) |
| pOFC | AntCing_MedPFC | L | 0.0040 | 0.9999*** | [0.9999, 0.9999] | (-14.0, 16.0, -18.0) |
| s32 | AntCing_MedPFC | L | 0.0330 | 0.2242 | [-0.2310, 0.5989] | (-10.0, 36.0, -10.0) |
| 11l | OrbPolaFrontal | L | 0.1266 | 0.8065* | [0.5740, 0.9186] | (-30.0, 54.0, -12.0) |
| 13l | OrbPolaFrontal | L | -0.0035 | 0.6454 | [0.2950, 0.8428] | (-24.0, 20.0, -24.0) |
| 47m | OrbPolaFrontal | L | -0.2705 | 0.5935 | [-0.2390, 0.5935] | (-28.0, 36.0, -8.0) |
| OFC | OrbPolaFrontal | L | -0.0407 | 0.9863*** | [0.9659, 0.9945] | (-18.0, 20.0, -22.0) |
| a47r | Inferior_Frontal | L | 0.1093 | 0.6723 | [0.3377, 0.8560] | (-34.0, 54.0, -12.0) |
| p47r | Inferior_Frontal | L | -0.0239 | 0.6414 | [0.2887, 0.8408] | (-52.0, 44.0, 8.0) |
| 10r | AntCing_MedPFC | R | 0.2780 | 0.7040 | [0.3901, 0.8713] | (4.0, 64.0, -4.0) |
| 10v | AntCing_MedPFC | R | 0.1445 | 0.9997*** | [0.9993, 0.9998] | (4.0, 68.0, -10.0) |
| 25 | AntCing_MedPFC | R | -0.0870 | 0.6499 | [0.3021, 0.8450] | (2.0, 16.0, -20.0) |
| a24 | AntCing_MedPFC | R | -0.0645 | 0.3388 | [-0.1101, 0.6730] | (8.0, 36.0, 6.0) |
| a32pr | AntCing_MedPFC | R | -0.1417 | 0.3868 | [-0.0552, 0.7021] | (16.0, 38.0, 20.0) |
| d32 | AntCing_MedPFC | R | -0.2009 | 0.2898 | [-0.1635, 0.6421] | (16.0, 40.0, 20.0) |
| pOFC | AntCing_MedPFC | R | 0.0508 | 0.9953*** | [0.9882, 0.9981] | (6.0, 12.0, -22.0) |
| s32 | AntCing_MedPFC | R | -0.0275 | 0.1290 | [-0.3217, 0.5321] | (4.0, 44.0, -12.0) |
| 11l | OrbPolaFrontal | R | 0.1400 | 0.9997*** | [0.9994, 0.9999] | (20.0, 58.0, -20.0) |
| 13l | OrbPolaFrontal | R | 0.1007 | 0.9874*** | [0.9687, 0.9950] | (20.0, 30.0, -24.0) |
| 47m | OrbPolaFrontal | R | 0.1346 | 0.5372 | [0.1360, 0.7870] | (30.0, 38.0, -6.0) |
| OFC | OrbPolaFrontal | R | -0.0168 | 0.9929*** | [0.9822, 0.9971] | (18.0, 30.0, -24.0) |
| a47r | Inferior_Frontal | R | 0.2929 | 0.5854 | [0.2044, 0.8124] | (42.0, 58.0, -6.0) |
| p47r | Inferior_Frontal | R | -0.0710 | 0.3179 | [-0.1332, 0.6600] | (50.0, 48.0, -6.0) |
| | | | Striatum | | | |
| Putam | Subcortical | L | -0.1602 | 0.5599 | [0.1677, 0.7991] | (-16.0, 10.0, -14.0) |
| Caud | Subcortical | L | -0.0038 | 0.4551 | [0.0277, 0.7418] | (-12.0, 24.0, -4.0) |
| NAc | Subcortical | L | -0.0646 | 0.6199 | [0.2557, 0.8300] | (-8.0, 8.0, -12.0) |
| Putam | Subcortical | R | -0.1739 | 0.3742 | [-0.0699, 0.6945] | (24.0, 6.0, 14.0) |
| Caud | Subcortical | R | 0.0972 | 0.7356 | [0.4443, 0.8863] | (26.0, -32.0, 12.0) |
| NAc | Subcortical | R | 0.1558 | 0.6057 | [0.2343, 0.8228] | (10.0, 10.0, -12.0) |

*For each a priori region of interest, the table summarizes the location of the most reliable voxel during stimulus processing, together with its peak intraclass correlation coefficient (ICC) and MNI coordinates for the delayed > immediate contrast. Asterisks indicate the reliability level of the peak ICC based on the lower bound of the 95% confidence interval: * moderate reliability (> 0.50), ** good reliability (> 0.75), *** excellent reliability (> 0.90).*

***Table S4. Peak ICC values of key region defined by the HCPex atlas during feedback presentation for the main effect contrast (delayed > immediate) using PyReliMRI's default method.***

| Parcel | Cortical Division | Side | Mean ICC | Peak ICC | 95% CI | Peak MNI (x, y, z) |
|---|---|---|---|---|---|---|
| | | | DLPFC | | | |
| 46 | Dorsolateral_Prefrontal | L | -0.0761 | 0.4881 | [0.0700, 0.7600] | (-32.0, 40.0, 30.0) |
| 8Ad | Dorsolateral_Prefrontal | L | -0.0025 | 0.4486 | [0.0200, 0.7380] | (-28.0, 20.0, 48.0) |
| 8Av | Dorsolateral_Prefrontal | L | 0.0515 | 0.9911*** | [0.9780, 0.9960] | (-40.0, 26.0, 52.0) |
| 8BL | Dorsolateral_Prefrontal | L | -0.0691 | 0.5345 | [0.1320, 0.7860] | (-8.0, 42.0, 50.0) |
| 8C | Dorsolateral_Prefrontal | L | -0.2450 | 0.4651 | [0.0400, 0.7470] | (-48.0, 24.0, 40.0) |
| 9-46d | Dorsolateral_Prefrontal | L | -0.0509 | 0.5240 | [0.1180, 0.7800] | (-26.0, 50.0, 6.0) |
| 9a | Dorsolateral_Prefrontal | L | -0.1098 | 0.5290 | [0.1250, 0.7830] | (-26.0, 54.0, 12.0) |
| 9p | Dorsolateral_Prefrontal | L | -0.1194 | 0.3744 | [-0.0700, 0.6949] | (-12.0, 50.0, 34.0) |
| a9-46v | Dorsolateral_Prefrontal | L | -0.1116 | 0.3941 | [-0.0470, 0.7060] | (-30.0, 54.0, 6.0) |
| i6-8 | Dorsolateral_Prefrontal | L | -0.0244 | 0.3913 | [-0.0500, 0.7050] | (-28.0, 10.0, 50.0) |
| s6-8 | Dorsolateral_Prefrontal | L | 0.0582 | 0.2934 | [-0.1600, 0.6440] | (-24.0, 22.0, 64.0) |
| SFL | Dorsolateral_Prefrontal | L | -0.2564 | 0.2133 | [-0.2420, 0.5920] | (-10.0, 20.0, 70.0) |
| | | | vmPFC/OFC and ACC/mPFC | | | |
| 10r | AntCing_MedPFC | L | -0.0645 | 0.3631 | [-0.0827, 0.6878] | (-12.0, 46.0, -10.0) |
| 10v | AntCing_MedPFC | L | -0.0729 | 0.9856*** | [0.9641, 0.9943] | (-6.0, 66.0, -8.0) |
| 25 | AntCing_MedPFC | L | -0.1307 | 0.5951 | [0.2185, 0.8174] | (-4.0, 14.0, -16.0) |
| a24 | AntCing_MedPFC | L | -0.1423 | 0.3412 | [-0.1074, 0.6745] | (-2.0, 50.0, -6.0) |
| a32pr | AntCing_MedPFC | L | -0.2946 | 0.0539 | [-0.3880, 0.4756] | (-8.0, 24.0, 36.0) |
| d32 | AntCing_MedPFC | L | -0.1148 | 0.2502 | [-0.2047, 0.6163] | (-8.0, 48.0, 34.0) |
| pOFC | AntCing_MedPFC | L | -0.2175 | 0.9998*** | [0.9997, 0.9999] | (-4.0, 12.0, -20.0) |
| s32 | AntCing_MedPFC | L | -0.1281 | 0.1031 | [-0.3451, 0.5130] | (-12.0, 40.0, -10.0) |
| 11l | OrbPolaFrontal | L | -0.1709 | 0.9821*** | [0.9555, 0.9928] | (-28.0, 66.0, -14.0) |
| 13l | OrbPolaFrontal | L | -0.1880 | 0.4209 | [-0.0145, 0.7222] | (-24.0, 20.0, -24.0) |
| 47m | OrbPolaFrontal | L | -0.1571 | 0.1997 | [-0.2551, 0.5822] | (-42.0, 32.0, -16.0) |
| OFC | OrbPolaFrontal | L | 0.0780 | 0.9999*** | [0.9999, 0.9999] | (-4.0, 34.0, -26.0) |
| a47r | Inferior_Frontal | L | 0.0401 | 0.4178 | [-0.0183, 0.7203] | (-38.0, 56.0, -10.0) |
| p47r | Inferior_Frontal | L | -0.1366 | 0.2325 | [-0.2227, 0.6045] | (-44.0, 40.0, -6.0) |
| 10r | AntCing_MedPFC | R | 0.2658 | 0.8562* | [0.6727, 0.9405] | (12.0, 50.0, -12.0) |
| 10v | AntCing_MedPFC | R | -0.0463 | 0.9677*** | [0.9204, 0.9873] | (4.0, 66.0, -16.0) |
| 25 | AntCing_MedPFC | R | -0.2344 | 0.7176 | [0.4131, 0.8778] | (2.0, 14.0, -20.0) |
| a24 | AntCing_MedPFC | R | -0.1402 | 0.3620 | [-0.0839, 0.6872] | (8.0, 36.0, 6.0) |
| a32pr | AntCing_MedPFC | R | -0.3617 | 0.1272 | [-0.3233, 0.5308] | (16.0, 38.0, 20.0) |
| d32 | AntCing_MedPFC | R | -0.1893 | 0.2005 | [-0.2543, 0.5828] | (4.0, 48.0, 28.0) |
| pOFC | AntCing_MedPFC | R | -0.0153 | 0.9861*** | [0.9652, 0.9944] | (14.0, 14.0, -20.0) |
| s32 | AntCing_MedPFC | R | -0.0798 | 0.0709 | [-0.3734, 0.4887] | (6.0, 44.0, -14.0) |
| 11l | OrbPolaFrontal | R | 0.0492 | 0.9422** | [0.8601, 0.9767] | (32.0, 64.0, -16.0) |
| 13l | OrbPolaFrontal | R | 0.0507 | 0.8484* | [0.6568, 0.9371] | (18.0, 20.0, -22.0) |
| 47m | OrbPolaFrontal | R | -0.0317 | 0.4569 | [0.0300, 0.7428] | (30.0, 38.0, -6.0) |
| OFC | OrbPolaFrontal | R | -0.0895 | 0.9995*** | [0.9988, 0.9998] | (16.0, 30.0, -24.0) |
| a47r | Inferior_Frontal | R | 0.1304 | 0.7073 | [0.3956, 0.8729] | (38.0, 64.0, 0.0) |
| p47r | Inferior_Frontal | R | -0.1025 | 0.2241 | [-0.2311, 0.5988] | (42.0, 42.0, -4.0) |
| | | | Striatum | | | |
| Putam | Subcortical | L | -0.0347 | 0.5341 | [0.1317, 0.7854] | (-28.0, 2.0, 0.0) |
| Caud | Subcortical | L | 0.0783 | 0.5808 | [0.1976, 0.8100] | (-16.0, -12.0, 26.0) |
| NAc | Subcortical | L | -0.4043 | -0.0364 | [-0.4620, 0.4027] | (-16.0, 8.0, -14.0) |
| Putam | Subcortical | R | -0.0788 | 0.5710 | [0.1836, 0.8049] | (22.0, 20.0, 2.0) |
| Caud | Subcortical | R | 0.2247 | 0.6390 | [0.2850, 0.8396] | (26.0, -32.0, 12.0) |
| NAc | Subcortical | R | -0.1186 | 0.1811 | [-0.2731, 0.5693] | (10.0, 8.0, -4.0) |

*For each a priori region of interest, the table summarizes the location of the most reliable voxel during feedback processing, together with its peak intraclass correlation coefficient (ICC) and MNI coordinates for the delayed > immediate contrast. Asterisks indicate the reliability level of the peak ICC based on the lower bound of the 95% confidence interval: * moderate reliability (> 0.50), ** good reliability (> 0.75), *** excellent reliability (> 0.90).*

TRR Contrast Results (Minimum Voxel Coverage Criterion) - Stimulus Presentation

As part of the main effect contrast analysis retaining only voxels with valid data from at least 10 participants, TRR of BOLD responses during stimulus presentation was assessed by computing ICC maps for the delayed > immediate contrast and overlaying them with the HCPex atlas.

Left dorsolateral prefrontal cortex (DLPFC) parcels showed poor-to-moderate peak ICC point estimates, ranging from 0.430 to 0.725. The highest peak reliability was observed in Area a9-46v (MNI: -48, 46, 14; ICC = 0.725), followed by Area 8Av (MNI: -32, 18, 58; ICC = 0.705), Area 8BL (MNI: -20, 30, 60; ICC = 0.697), and Area 8C (MNI: -48, 20, 44; ICC = 0.674). Area 46 also showed moderate peak reliability (MNI: -46, 44, 22; ICC = 0.623). However, none of the left DLPFC parcels had a 95% confidence interval lower bound exceeding 0.50.

ACC/mPFC parcels showed poor-to-moderate peak ICC estimates, ranging from 0.224 to 0.661. The highest value within this division was observed in right Area 10r (MNI: 12, 50, -12; ICC = 0.661). vmPFC/OFC parcels, together with adjacent inferior frontal parcels, showed more variable reliability, with peak ICC estimates ranging from 0.216 to 0.806. The highest peak reliability was observed in left Area 11l (MNI: -30, 54, -12; ICC = 0.806), followed by left Area a47r (MNI: -34, 54, -12; ICC = 0.672), and left Area p47r (MNI: -52, 44, 8; ICC = 0.641). Among these cortical parcels, only left Area 11l had a 95% confidence interval lower bound exceeding 0.50.

Striatal parcels showed poor-to-moderate peak ICC estimates, ranging from 0.196 to 0.736. In the right hemisphere, caudate showed the highest striatal point estimate peak reliability (MNI: 26, -32, 12; ICC = 0.735), followed by nucleus accumbens (MNI: 12, 14, -10; ICC = 0.573). In the left hemisphere, caudate showed poor reliability (MNI: -12, 24, -4; ICC = 0.455), whereas putamen and nucleus accumbens showed lower peak ICC estimates. No striatal parcel had a 95% confidence interval lower bound exceeding 0.50. Full stimulus-related contrast results are reported in Table S5.

TRR Contrast Results (Minimum Voxel Coverage Criterion) - Feedback Presentation

As part of the main effect contrast analysis retaining only voxels with valid data from at least 10 participants, TRR of BOLD responses during feedback presentation was assessed by computing ICC maps for the delayed > immediate contrast and overlaying them with the HCPex atlas.

Left DLPFC parcels showed predominantly poor-to-moderate peak ICC estimates, ranging from 0.213 to 0.534. The highest reliability was observed in Area 8BL (MNI: -8, 42, 50; ICC = 0.534), followed by Area 9a (MNI: -26, 54, 12; ICC = 0.529) and Area 9-46d (MNI: -26, 50, 6; ICC = 0.524). Area 46 showed a peak ICC estimate of 0.488 (MNI: -32, 40, 30). None of the left DLPFC parcels had a 95% confidence interval lower bound exceeding 0.50.

ACC/mPFC parcels showed generally poor peak point estimate reliability during feedback presentation, except for right Area 10r (MNI: 12, 50, -12; ICC = 0.856) and Area 10v (MNI: 6, 50, -12; ICC = 0.528). vmPFC/OFC, together with adjacent inferior frontal parcels, showed variable peak ICC estimates, ranging from -0.004 to 0.707. The strongest peak point estimate reliability was observed in right Area a47r (MNI: 38, 64, 0; ICC = 0.707) and right Area 11l (MNI: 18, 46, -16; ICC = 0.643). Of these parcels, only right Area 10r had a 95% confidence interval lower bound exceeding 0.50.

Striatal parcels showed variable peak ICC estimates, ranging from -0.128 to 0.639. In the right hemisphere, caudate showed the highest striatal reliability (MNI: 26, -32, 12; ICC = 0.639), followed by putamen (MNI: 22, 20, 2; ICC = 0.571). In the left hemisphere, caudate (MNI: -16, -12, 26; ICC = 0.581) and putamen (MNI: -28, 2, 0; ICC = 0.534) showed moderate peak reliability, whereas nucleus accumbens showed negative reliability (MNI: -8, 10, -4; ICC = -0.128). No striatal parcel had a 95% confidence interval lower bound exceeding 0.50. Full feedback-related contrast results are reported in Table S6.

***Figure S3. Whole-brain voxel-wise heat map of TRR across task sessions for BOLD responses of the main effect contrast (delayed > immediate). With minimum Voxel Coverage Criterion applied.***

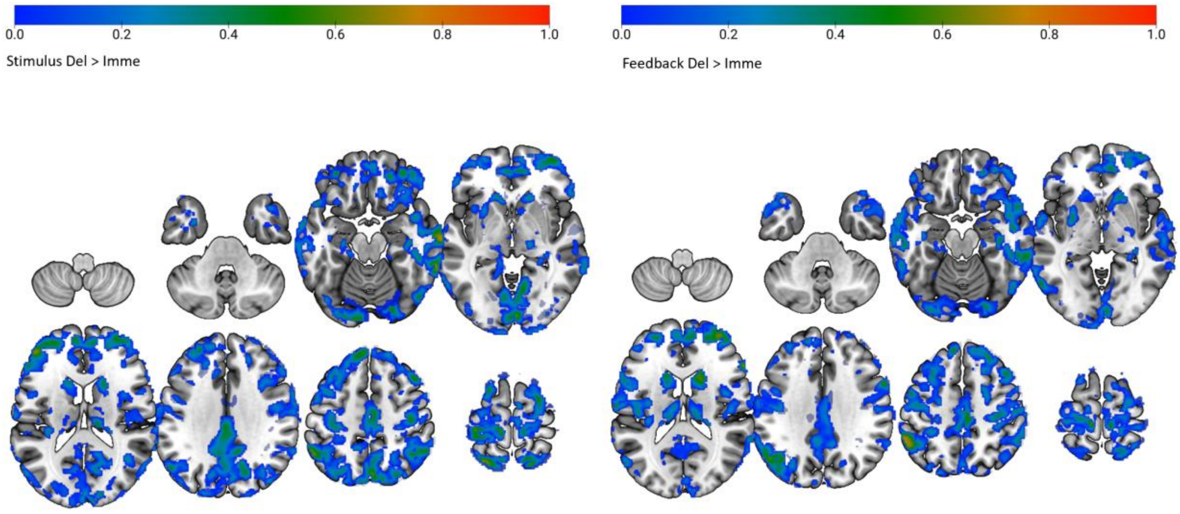


*Whole-brain voxel-wise ICC map computed for Delayed > Immediate for stimulus-related activity (left) and feedback-related activity (right) across the two task sessions.*

***Table S5. Peak ICC values of key region defined by the HCPex atlas during stimulus presentation for the main effect contrast (delayed > immediate) using Minimum Voxel Coverage Criterion.***

| Parcel | Cortical Division | Side | Mean ICC | Peak ICC | 95% CI | Peak MNI (x, y, z) |
|---|---|---|---|---|---|---|
| | | | DLPFC | | | |
| 46 | Dorsolateral_Prefrontal | L | 0.0127 | 0.6229 | [0.2602, 0.8316] | (-46.0, 44.0, 22.0) |
| 8Ad | Dorsolateral_Prefrontal | L | 0.1572 | 0.4490 | [0.0200, 0.7384] | (-16.0, 34.0, 38.0) |
| 8Av | Dorsolateral_Prefrontal | L | 0.2017 | 0.7059 | [0.3933, 0.8723] | (-32.0, 18.0, 58.0) |
| 8BL | Dorsolateral_Prefrontal | L | 0.1678 | 0.6977 | [0.3795, 0.8683] | (-20.0, 30.0, 60.0) |
| 8C | Dorsolateral_Prefrontal | L | -0.1134 | 0.6747 | [0.3417, 0.8572] | (-48.0, 20.0, 44.0) |
| 9-46d | Dorsolateral_Prefrontal | L | 0.0720 | 0.5750 | [0.1893, 0.8070] | (-24.0, 54.0, 12.0) |
| 9a | Dorsolateral_Prefrontal | L | 0.0648 | 0.6076 | [0.2371, 0.8238] | (-28.0, 56.0, 12.0) |
| 9p | Dorsolateral_Prefrontal | L | 0.0939 | 0.4636 | [0.0384, 0.7466] | (-24.0, 48.0, 34.0) |
| a9-46v | Dorsolateral_Prefrontal | L | 0.2677 | 0.7255 | [0.4265, 0.8815] | (-48.0, 46.0, 14.0) |
| i6-8 | Dorsolateral_Prefrontal | L | -0.0138 | 0.4305 | [-0.0029, 0.7277] | (-40.0, 2.0, 62.0) |
| s6-8 | Dorsolateral_Prefrontal | L | 0.1249 | 0.6614 | [0.3202, 0.8507] | (-22.0, 26.0, 62.0) |
| SFL | Dorsolateral_Prefrontal | L | -0.3294 | 0.4709 | [0.0477, 0.7507] | (-8.0, 24.0, 68.0) |
| | | | vmPFC/OFC and ACC/mPFC | | | |
| 10r | AntCing_MedPFC | L | 0.0653 | 0.5047 | [0.0919, 0.7694] | (-10.0, 48.0, -8.0) |
| 10v | AntCing_MedPFC | L | -0.0411 | 0.3843 | [-0.0583, 0.7006] | (-6.0, 58.0, -16.0) |
| 25 | AntCing_MedPFC | L | -0.1584 | 0.3319 | [-0.1179, 0.6687] | (0.0, 10.0, -10.0) |
| a24 | AntCing_MedPFC | L | -0.0487 | 0.5093 | [0.0981, 0.7720] | (-2.0, 50.0, -6.0) |
| a32pr | AntCing_MedPFC | L | -0.1092 | 0.2494 | [-0.2056, 0.6158] | (-10.0, 26.0, 34.0) |
| d32 | AntCing_MedPFC | L | -0.1083 | 0.3617 | [-0.0844, 0.6870] | (-14.0, 48.0, 10.0) |
| pOFC | AntCing_MedPFC | L | -0.0471 | 0.2221 | [-0.2331, 0.5975] | (-14.0, 16.0, -14.0) |
| s32 | AntCing_MedPFC | L | 0.0331 | 0.2242 | [-0.2311, 0.5989] | (-10.0, 36.0, -10.0) |
| 11l | OrbPolaFrontal | L | 0.2571 | 0.8065* | [0.5741, 0.9187] | (-30.0, 54.0, -12.0) |
| 13l | OrbPolaFrontal | L | -0.0678 | 0.4605 | [0.0346, 0.7449] | (-22.0, 38.0, -12.0) |
| 47m | OrbPolaFrontal | L | -0.2706 | 0.2162 | [-0.2390, 0.5935] | (-28.0, 36.0, -8.0) |
| OFC | OrbPolaFrontal | L | 0.0281 | 0.6101 | [0.2409, 0.8251] | (-12.0, 36.0, -20.0) |
| a47r | Inferior_Frontal | L | 0.1120 | 0.6723 | [0.3377, 0.8560] | (-34.0, 54.0, -12.0) |
| p47r | Inferior_Frontal | L | -0.0251 | 0.6415 | [0.2888, 0.8408] | (-52.0, 44.0, 8.0) |
| 10r | AntCing_MedPFC | R | 0.2755 | 0.6618 | [0.3208, 0.8509] | (12.0, 50.0, -12.0) |
| 10v | AntCing_MedPFC | R | 0.0693 | 0.4807 | [0.0604, 0.7562] | (6.0, 50.0, -16.0) |
| 25 | AntCing_MedPFC | R | -0.1179 | 0.6407 | [0.2875, 0.8404] | (6.0, 12.0, -12.0) |
| a24 | AntCing_MedPFC | R | -0.0645 | 0.3389 | [-0.1101, 0.6730] | (8.0, 36.0, 6.0) |
| a32pr | AntCing_MedPFC | R | -0.1417 | 0.3869 | [-0.0552, 0.7021] | (16.0, 38.0, 20.0) |
| d32 | AntCing_MedPFC | R | -0.2010 | 0.2899 | [-0.1635, 0.6421] | (16.0, 40.0, 20.0) |
| pOFC | AntCing_MedPFC | R | -0.0227 | 0.6059 | [0.2345, 0.8229] | (12.0, 16.0, -12.0) |
| s32 | AntCing_MedPFC | R | -0.0276 | 0.1291 | [-0.3218, 0.5322] | (4.0, 44.0, -12.0) |
| 11l | OrbPolaFrontal | R | 0.1845 | 0.6061 | [0.2349, 0.8230] | (36.0, 54.0, -10.0) |
| 13l | OrbPolaFrontal | R | 0.0301 | 0.3927 | [-0.0484, 0.7056] | (20.0, 42.0, -18.0) |
| 47m | OrbPolaFrontal | R | 0.1347 | 0.5373 | [0.1361, 0.7871] | (30.0, 38.0, -6.0) |
| OFC | OrbPolaFrontal | R | 0.0253 | 0.3449 | [-0.1034, 0.6767] | (8.0, 32.0, -22.0) |
| a47r | Inferior_Frontal | R | 0.3014 | 0.5855 | [0.2045, 0.8124] | (42.0, 58.0, -6.0) |
| p47r | Inferior_Frontal | R | -0.0711 | 0.3180 | [-0.1332, 0.6600] | (50.0, 48.0, -6.0) |
| | | | Striatum | | | |
| Putam | Subcortical | L | -0.1591 | 0.2707 | [-0.1836, 0.6298] | (-26.0, 10.0, -2.0) |
| Caud | Subcortical | L | -0.0038 | 0.4552 | [0.0278, 0.7419] | (-12.0, 24.0, -4.0) |
| NAc | Subcortical | L | -0.0916 | 0.1961 | [-0.2587, 0.5797] | (-6.0, 10.0, -10.0) |
| Putam | Subcortical | R | -0.1739 | 0.3742 | [-0.0700, 0.6946] | (24.0, 6.0, 14.0) |
| Caud | Subcortical | R | 0.0972 | 0.7357 | [0.4443, 0.8863] | (26.0, -32.0, 12.0) |
| NAc | Subcortical | R | 0.1620 | 0.5729 | [0.1862, 0.8059] | (12.0, 14.0, -10.0) |

*For each a priori region of interest, the table summarizes the location of the most reliable voxel during stimulus processing, together with its peak intraclass correlation coefficient (ICC) and MNI coordinates for the delayed > immediate contrast. Asterisks indicate the reliability level of the peak ICC based on the lower bound of the 95% confidence interval: * moderate reliability (> 0.50), ** good reliability (> 0.75), *** excellent reliability (> 0.90).*

***Table S6. Peak ICC values of key region defined by the HCPex atlas during feedback presentation for the main effect contrast (delayed > immediate) using Minimum Voxel Coverage Criterion.***

| Parcel | Cortical Division | Side | Mean ICC | Peak ICC | 95% CI | Peak MNI (x, y, z) |
|---|---|---|---|---|---|---|
| | | | DLPFC | | | |
| 46 | Dorsolateral_Prefrontal | L | -0.0594 | 0.4881 | [0.0700, 0.7603] | (-32.0, 40.0, 30.0) |
| 8Ad | Dorsolateral_Prefrontal | L | -0.0046 | 0.4486 | [0.0196, 0.7382] | (-28.0, 20.0, 48.0) |
| 8Av | Dorsolateral_Prefrontal | L | 0.0372 | 0.4996 | [0.0852, 0.7667] | (-30.0, 18.0, 58.0) |
| 8BL | Dorsolateral_Prefrontal | L | -0.0732 | 0.5345 | [0.1322, 0.7856] | (-8.0, 42.0, 50.0) |
| 8C | Dorsolateral_Prefrontal | L | -0.2475 | 0.4651 | [0.0404, 0.7475] | (-48.0, 24.0, 40.0) |
| 9-46d | Dorsolateral_Prefrontal | L | -0.0383 | 0.5240 | [0.1179, 0.7799] | (-26.0, 50.0, 6.0) |
| 9a | Dorsolateral_Prefrontal | L | -0.0912 | 0.5290 | [0.1247, 0.7826] | (-26.0, 54.0, 12.0) |
| 9p | Dorsolateral_Prefrontal | L | -0.1002 | 0.3744 | [-0.0698, 0.6947] | (-12.0, 50.0, 34.0) |
| a9-46v | Dorsolateral_Prefrontal | L | -0.1091 | 0.3941 | [-0.0467, 0.7064] | (-30.0, 54.0, 6.0) |
| i6-8 | Dorsolateral_Prefrontal | L | -0.0192 | 0.3913 | [-0.0501, 0.7048] | (-28.0, 10.0, 50.0) |
| s6-8 | Dorsolateral_Prefrontal | L | -0.0572 | 0.2599 | [-0.1949, 0.6227] | (-22.0, 22.0, 50.0) |
| SFL | Dorsolateral_Prefrontal | L | -0.2562 | 0.2133 | [-0.2418, 0.5916] | (-10.0, 20.0, 70.0) |
| | | | vmPFC/OFC and ACC/mPFC | | | |
| 10r | AntCing_MedPFC | L | -0.0211 | 0.3631 | [-0.0828, 0.6879] | (-12.0, 46.0, -10.0) |
| 10v | AntCing_MedPFC | L | -0.1720 | 0.1291 | [-0.3217, 0.5322] | (0.0, 36.0, -22.0) |
| 25 | AntCing_MedPFC | L | -0.1430 | 0.2734 | [-0.1808, 0.6315] | (-6.0, 14.0, -14.0) |
| a24 | AntCing_MedPFC | L | -0.1423 | 0.3413 | [-0.1074, 0.6745] | (-2.0, 50.0, -6.0) |
| a32pr | AntCing_MedPFC | L | -0.2947 | 0.0539 | [-0.3880, 0.4757] | (-8.0, 24.0, 36.0) |
| d32 | AntCing_MedPFC | L | -0.1148 | 0.2503 | [-0.2048, 0.6164] | (-8.0, 48.0, 34.0) |
| pOFC | AntCing_MedPFC | L | -0.2433 | -0.0046 | [-0.4366, 0.4291] | (-12.0, 22.0, -18.0) |
| s32 | AntCing_MedPFC | L | -0.1281 | 0.1031 | [-0.3452, 0.5131] | (-12.0, 40.0, -10.0) |
| 11l | OrbPolaFrontal | L | -0.0810 | 0.5355 | [0.1336, 0.7861] | (-30.0, 54.0, -14.0) |
| 13l | OrbPolaFrontal | L | -0.2607 | 0.2565 | [-0.1984, 0.6205] | (-18.0, 32.0, -20.0) |
| 47m | OrbPolaFrontal | L | -0.1572 | 0.1998 | [-0.2551, 0.5823] | (-42.0, 32.0, -16.0) |
| OFC | OrbPolaFrontal | L | -0.1617 | 0.3366 | [-0.1127, 0.6716] | (-12.0, 38.0, -20.0) |
| a47r | Inferior_Frontal | L | 0.0396 | 0.4178 | [-0.0183, 0.7204] | (-38.0, 56.0, -10.0) |
| p47r | Inferior_Frontal | L | -0.1378 | 0.2325 | [-0.2227, 0.6045] | (-44.0, 40.0, -6.0) |
| 10r | AntCing_MedPFC | R | 0.2893 | 0.8563* | [0.6728, 0.9405] | (12.0, 50.0, -12.0) |
| 10v | AntCing_MedPFC | R | -0.0643 | 0.5288 | [0.1245, 0.7826] | (6.0, 50.0, -16.0) |
| 25 | AntCing_MedPFC | R | -0.2465 | 0.1936 | [-0.2612, 0.5780] | (4.0, 14.0, -14.0) |
| a24 | AntCing_MedPFC | R | -0.1402 | 0.3621 | [-0.0839, 0.6872] | (8.0, 36.0, 6.0) |
| a32pr | AntCing_MedPFC | R | -0.3618 | 0.1273 | [-0.3234, 0.5309] | (16.0, 38.0, 20.0) |
| d32 | AntCing_MedPFC | R | -0.1894 | 0.2006 | [-0.2544, 0.5828] | (4.0, 48.0, 28.0) |
| pOFC | AntCing_MedPFC | R | -0.1142 | 0.4760 | [0.0544, 0.7536] | (24.0, 16.0, -22.0) |
| s32 | AntCing_MedPFC | R | -0.0799 | 0.0709 | [-0.3734, 0.4888] | (6.0, 44.0, -14.0) |
| 11l | OrbPolaFrontal | R | 0.0342 | 0.6435 | [0.2919, 0.8418] | (18.0, 46.0, -16.0) |
| 13l | OrbPolaFrontal | R | -0.0135 | 0.2369 | [-0.2184, 0.6075] | (24.0, 34.0, -10.0) |
| 47m | OrbPolaFrontal | R | -0.0318 | 0.4569 | [0.0300, 0.7429] | (30.0, 38.0, -6.0) |
| OFC | OrbPolaFrontal | R | -0.0681 | 0.2229 | [-0.2324, 0.5981] | (16.0, 32.0, -20.0) |
| a47r | Inferior_Frontal | R | 0.1343 | 0.7074 | [0.3956, 0.8729] | (38.0, 64.0, 0.0) |
| p47r | Inferior_Frontal | R | -0.1026 | 0.2241 | [-0.2311, 0.5989] | (42.0, 42.0, -4.0) |
| | | | Striatum | | | |
| Putam | Subcortical | L | -0.0329 | 0.5341 | [0.1317, 0.7854] | (-28.0, 2.0, 0.0) |
| Caud | Subcortical | L | 0.0784 | 0.5808 | [0.1977, 0.8100] | (-16.0, -12.0, 26.0) |
| NAc | Subcortical | L | -0.3347 | -0.1280 | [-0.5314, 0.3227] | (-8.0, 10.0, -4.0) |
| Putam | Subcortical | R | -0.0788 | 0.5711 | [0.1836, 0.8050] | (22.0, 20.0, 2.0) |
| Caud | Subcortical | R | 0.2248 | 0.6391 | [0.2851, 0.8397] | (26.0, -32.0, 12.0) |
| NAc | Subcortical | R | -0.0838 | 0.1812 | [-0.2731, 0.5694] | (10.0, 8.0, -4.0) |

*For each a priori region of interest, the table summarizes the location of the most reliable voxel during feedback processing, together with its peak intraclass correlation coefficient (ICC) and MNI coordinates for the delayed > immediate contrast. Asterisks indicate the reliability level of the peak ICC based on the lower bound of the 95% confidence interval: * moderate reliability (> 0.50), ** good reliability (> 0.75), *** excellent reliability (> 0.90).*
